# Monitor individual health and improve breeding success in Crested ibis (*Nipponia nippon*)

**DOI:** 10.64898/2025.12.02.691777

**Authors:** Yuansi He, Xu Zhang, Xuebo Xi, Shuai Yang, Dejing Cai, Xiangjiang Zhan, Daiping Wang

**Affiliations:** Institute of Zoology, Chinese Academy of Sciences; Administration Bureau of Dongzhai National Nature Reserve, Luoshan, Henan Province, China; Chinese Academy of Sciences

**Keywords:** Crested ibis (*Nipponia nippon*), Heritability, Breeding values, Health predicting

## Abstract

Biodiversity loss has become a pressing issue, requiring effective conservation measures. Drawing lessons from successful examples is essential. The Crested ibis (*Nipponia nippon*), once critically endangered but now recovering through intensive conservation programs, provides an informative model for evaluating and improving conservation practices. Using 17 years of monitoring data from a captive population spanning eight generations, we applied quantitative genetic tools (Animal Model) to characterize individual growth and improve breeding success of this endangered species. We found that body weight from Day 0 to Day 42 exhibited significant heritability (*h*^2^ = 0.195, 95% HPD 0.139 ∼ 0.250). As for growth curve parameters, growth time-scale parameter ***a*** (*h^2^* = 0.33, 95% HPD: 0.14 ∼ 0.55) and growth-rate coefficient ***b*** (*h*^2^ = 0.22, 95% HPD: 0.10 ∼ 0.34) were also significantly heritable, while the heritability of potential maximum weight ***M*** was relatively low (*h^2^* = 0.023, 95% HPD: 0.00 ∼ 0.16). We further estimated the breeding values of these phenotypical traits to inform future breeding and selections. Intergenerational analyses showed the estimated breeding values for body weight exhibited a tendency to increase, accompanied by a slower growth rate, though trends did not differ significantly from expectations under genetic drift. Comparison of parental and offspring growth trajectories revealed that healthy offspring closely follow parental growth trajectories, whereas unhealthy individuals display reduced growth. Given the multiple breeding times of this long-lived species, this approach enables effective monitoring of individual growth and health, allowing timely veterinary interventions so to enhance conservation efficiency.

## Introduction

Global biodiversity is being degraded at an unprecedented rate. More than one in four species assessed by the IUCN is threatened with extinction (IUCN 2025), and this decline is accelerating under increasing pressures such as climate change, habitat loss and human disturbance (Krauss et al. 2010; Urban 2015; Isbell et al. 2023). Reversing this declining trend is an urgent global priority (Cowie et al. 2022). Effective conservation of rare and endangered species is therefore essential not only because they are in endangered status and require urgent protection, but also because many serve as flagship or umbrella species that enhance public engagement and promote the protection of broader ecosystems (Lambeck 1997; Zacharias and Roff 2001; Roberge and Angelstam 2004). At the same time, many conservation programs still show generally low long-term success rates (Seddon et al. 2014), underscoring the need to extract practical lessons from well-documented successful cases.

The Crested ibis (*Nipponia nippon*) is one of the most famous and successful examples of how conservation efforts have brought a species back from the brink of extinction. It was historically widespread in Northeast Asia (BirdLife-International 2001; Li et al. 2014), but its population experienced a dramatic decline in the 20^th^ century due to the destruction of habitats and human activity (e.g., illegal hunting, overuse of fertilizer and pesticide) (Archibald et al. 1980; Ding 2004; Li et al. 2009; Feng et al. 2019). Populations in Russia, Korea, and Japan disappeared in succession and the species was once presumed extinct in the wild (Archibald and Lantis 1979; Anderson 1984). The last small wild population rediscovered in Yangxian County of Shaanxi Province, China in 1981 contained only seven individuals, including two breeding pairs and three chicks from one of the couples (Liu 1981). After that, with extensive in-situ and ex-situ conservation efforts, the global population size of wild and captive Crested ibis has now been estimated to exceed 10,000 (Li 2023), with several populations located in China (Yu et al. 2015; Zheng et al. 2018; Xie et al. 2020; Qiu et al. 2023; Cai et al. 2024), Korea (Choi et al. 2020), and Japan (Wajiki et al. 2014; Okahisa et al. 2022). However, the species still suffers from inbreeding depression and low genetic diversity due to the extremely small number of founders, leading to high rates of embryonic mortality and congenital defects (Fu et al. 2019; Zheng et al. 2024), which is also a challenge that is widely shared across the conservation programs of many other species (Ballou et al. 1989; Beissinger et al. 2008; Grueber et al. 2010; Khan et al. 2021). Thus, further investigation is needed to detect the abnormality and reduce the negative impact. Despite remarkable progress in population recovery, including long-term successful captive breeding, previous studies about captive population in Crested ibis were mainly descriptive, focusing on physiological characteristics or feeding methods (Wingfield et al. 2000; Huang et al. 2006, 2016; Okahisa et al. 2022). Quantitative genetic parameters such as heritability and estimated breeding values for key traits remain poorly understood. This gap limits our ability to identify heritable traits, predict offspring performance from parental traits, detect individuals deviating from expected developmental patterns so that timely supportive interventions can be implemented, and evaluate population quality for the preservation of high-value genetic resources.

Verifying fitness-relevant traits of interest that are heritable and identifying the genetic differences behind them are important steps to improve the quality and breeding success of populations (Toghiani 2012). Quantitative genetic tools, especially the “Animal Model”, provide a powerful framework for partitioning phenotypic variation into genetic and environmental components using pedigree information. It enables estimation of heritability and individual breeding values, and further identification of superior individuals as well as preservation of valuable germplasm resources (Kruuk 2004; Wilson et al. 2010). These approaches are widely used in animal breeding to improve traits of economic interest, such as ranking individuals by their estimated breeding values of key traits to guide future breeding programs (Narinç et al. 2016; Abou Khadiga et al. 2016). It is also used in health-related trait selection to facilitate genetic improvement, such as selective breeding for immune-related traits (Mallard et al. 1992) or against hereditary defects (Malm et al. 2008). Beyond captive breeding, breeding values estimation has also been applied in research on natural populations and wildlife conservation, offering insights into natural selection and adaptive evolution, informing population management strategies, and enhancing the ability to predict population dynamics. It has been used to explain cases how evolutionary responses of heritable traits to strong and consistent directional selection are masked by environmental changes (Garant et al. 2004) or sexually antagonistic selection (Regan et al. 2019). Additionally, it has played an important role in health prediction and identification of genetic resources in endangered species (Guhlin et al. 2023). Overall, applying these methods in conservation programs can facilitate more efficient outcomes and enhance the quality of the targeted populations.

Body weight is a key fitness-related trait associated with survival rates (Krementz et al. 1989; Garant et al. 2004; Wilson et al. 2007; Tarwater et al. 2011; Ronget et al. 2018), reproductive success (Monaghan et al. 1989; Wendeln and Becker 1999), foraging ability (Donnelly and Sullivan 1998) and anti-predation capacity (Naef-Daenzer and Grüebler 2008). It is not only closely related to the current fitness, but also infect individual’s future survival (Merilä and Svensson 1997). It has been shown to be heritable in many species (Garant et al. 2004; Wilson et al. 2007; Saunders and Cuthbert 2014; Manjula et al. 2018; Singh et al. 2018), enabling predictions of offspring performance from parental phenotypes and facilitating selective breeding based on estimated breeding values. It also reflects environmental and nutritional conditions (Monaghan et al. 1989; Killpack and Karasov 2012), making it a useful indicator of health status. While body weight offers a snapshot of an individual’s condition at a given time, the growth trend during the nestling period offers a more comprehensive assessment of physiological condition and nutritional status (Boag 1987; Negro et al. 1994; N’Dri et al. 2006), and growth curve parameters have also been demonstrated to be heritable (Tholon et al. 2006; Narinc et al. 2014; Narinç et al. 2017). Therefore, these traits represent promising candidates for health assessment and selective breeding in the Crested ibis.

In this study, we focus on the body weight and Gompertz curve-fitted growth trend based on 17 years of accumulated data from a captive Crested ibis population. Our aim is to integrate quantitative genetics into conservation, which is the first time for this endangered species, assessing whether these essential fitness-related traits are heritable in Crested ibis and whether they can be used for further conservation improvement like health prediction and preservation of high-quality genetic resources. Specifically, we (1) estimated the heritability and breeding values of body weight and growth curve parameters; (2) assessed generational changes in estimated breeding values and compared the observed patterns with those expected under simulated genetic drift; and (3) evaluated parent-offspring differences in growth trajectories to identify unhealthy individuals and determine the timing for necessary veterinary interventions. We found that body weight and growth curve parameters, except for potential maximum weight, exhibited significant heritability, supporting their utility for selective breeding and health assessment. The body weight showed an overall increasing trend accompanied by slower growth rates across eight generations, though the observed changes did not differ significantly from drift expectations. Additionally, unhealthy offspring displayed poorer growth with unusual body weight inflection points that deviated from parental expectations, suggesting early developmental deviations can serve as practical indicators for targeted veterinary care. Overall, our study provides the first pedigree-based quantitative genetic analyses of fitness-related growth traits in the endangered Crested ibis, offering an evidence-based framework for predicting the growth trend, monitoring health, and improving conservation breeding outcomes. The analytical approach and results can also be applied to other endangered species, including long-term monitored wild populations with pedigree and phenotypic data.

## Material and methods

### Study species

The Crested ibis is a medium-sized wading bird with red facial skin and legs. Female and male birds are similar in appearance (i.e., sexual monomorphism) (Figure 1). Adult birds are generally white with orangish and cinnamon tones in tail, abdomen and flight feathers during the non-breeding seasons. While in breeding seasons, they secrete a black substance from a patch of skin in the neck and throat region, and apply this substance to their crest, scapulars, mantle and wings through ‘daubing behavior’, which usually occurs after bathing (Uchida 1970). It typically reaches sexual maturity at around 3 years old (ranging from 2 to 4 years old). The average age of first reproduction for females is 3.64 ± 1.36 years, while males reproduce slightly later, at 4.17 ± 1.47 years (Yu et al. 2010), although some captive individuals begin breeding as early as two years old and may even lay eggs at one year old. The species is long-lived, with most individuals living more than 20 years and the oldest recorded captive bird reaching 39 years (China National Radio 2025). The breeding season usually spans from late February to late June. Nest building, incubation, and chick provisioning are jointly undertaken by both parents (Zhai et al. 2011). Mating and nest-building occur from late February to early March, with egg-laying typically starting in mid-March to early April (Li and Huang 1986; Shi and Yu 1989; Wang and Shi 1999). The Crested ibis usually lays one clutch per year, but may produce a replacement clutch if the first one fails early in the season. Each clutch contains 2 ∼ 4 eggs, laid at one- or two-day intervals (Shi et al. 1999). In captivity, egg collection can stimulate additional laying, causing the total number of eggs to exceed the natural level, sometimes reaching more than ten. The incubation period lasts around 28 days, and chicks remain in the nest for about 40 ∼ 45 days before fledging (Shi et al. 1999; Zhai et al. 1999). After fledging, young birds continue to follow their parents and forage near the nesting tree (Ding et al. 1999).

**Figure 1.**
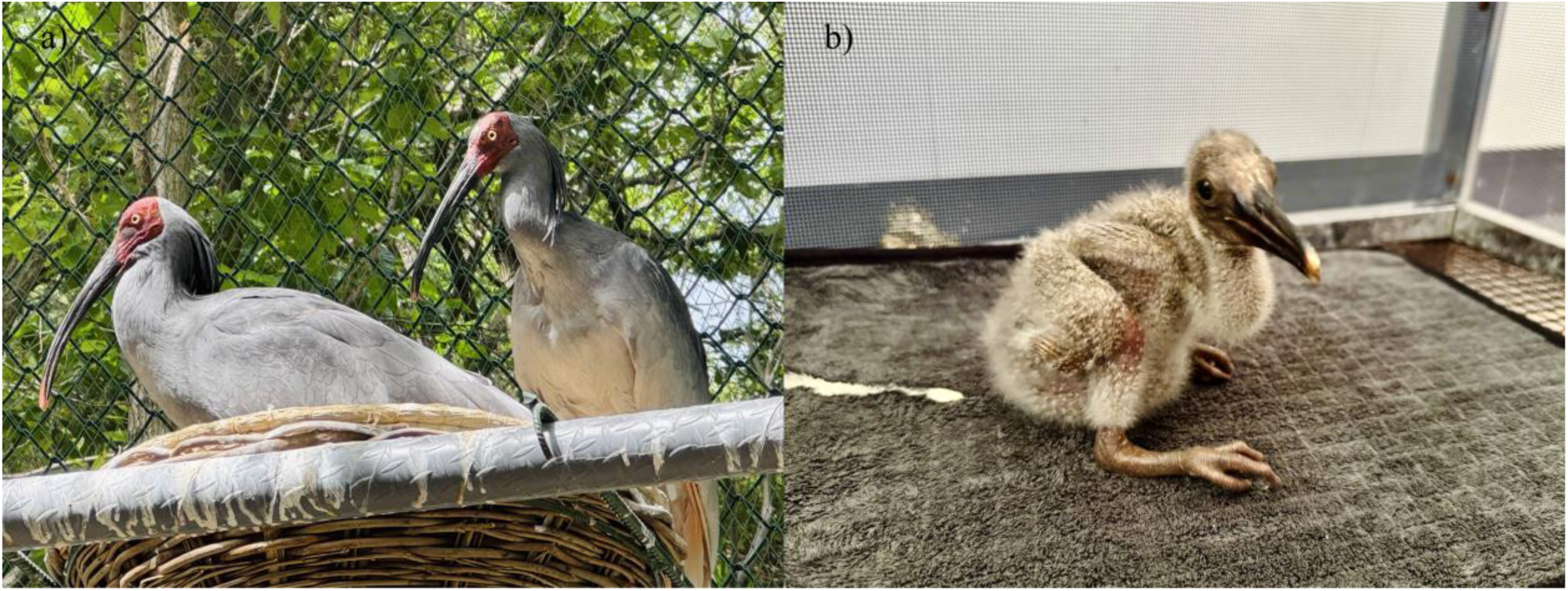
A pair of Crested ibis during the breeding season, with the male on the left and the female on the right (a), and a 9-day-old juvenile chick in artificial feeding environment (b); Photo by Yuansi He.

### Source of data

The dataset used in this study was collected from the Crested Ibis Breeding Center at Dongzhai National Nature Reserve, Henan Province, China. The reserve is located at E 114°18’ ∼ 114° 30’ and N 31°28’ ∼ 32°09’. The reintroduction program in this reserve started in 2007. A total of 17 founders were transferred to the breeding center, consisting of 4 individuals from the Beijing Zoo and 13 individuals from Japan (unfortunately, 4 of these founders died without successful breeding). It was the first time Crested ibis was reintroduced to its historical distribution outside Shaanxi Province in China. From 2008 to 2025, a total of 1,040 eggs were laid and 423 individuals were bred in captivity. Among these, 39 individuals had their parents or ancestors with unknown identities. The dataset includes each individual’s birth year, daily body weight, health status, and pedigree information. The pedigree structure of the captive population (eight generations) was presented in Figure S1 and was constructed using the package **pedantics** (Morrissey 2018). The founders were designated as Generation 1 and their parents were assigned to Generation 0. Subsequent generations were categorized according to the lower generational rank of their parents (e.g., if an individual’s father belonged to Generation 2 and the mother to Generation 3, then it would be placed in Generation 4).

### Traits measuring and management

#### 1) Body weight

Of all the 384 captive-bred individuals, we excluded those with less than 5 recorded measurements from 0 to 42 days old, considering that most nestlings fledged at around 40 to 45 days. At last, our dataset included 6422 daily weight records for 264 individuals born in captivity from 2008 to 2025. Of these, 102 were males, 97 were females and 65 remained unknown in gender. As for health status, 243 were healthy ones and 21 were unhealthy. An unhealthy individual was defined as one who died before reaching sexual maturity due to non-accidental causes, or had been clearly documented to have severe disabilities such as developmental delays, inability to walk, or eye diseases. All but two of them were hand-hatched in the incubator, with the parents initially incubating the eggs for 0 to about 25 days. Two chicks were hatched by their parents and later transitioned to artificial rearing due to the lack of feeding from the adult birds. All hatchlings were hand-reared by dedicated staffs under standardized conditions until they could feed themselves. Thus, unlike the wild populations, variation in growth was not influenced by nest differences or parental investment. Chicks were weighed daily when possible.

#### 2) Growth trend modeled by Gompertz curve

Chick growth was modeled by a three-parameter Gompertz curve, which has been widely used in animal studies. The Gompertz curve fits the potential maximum weight as parameter ***M***, the growth time-scale parameter as ***a*** and the growth-rate coefficient as ***b*** (Winsor 1932; Guhlin et al. 2023). Following the method of (Guhlin et al. 2023), the assumed model was:

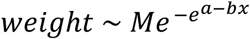

where ***weight*** was the body weight of an individual at ***x*** days old; ***M*** was the potential maximum weight the individual could reach; ***a*** was the growth time-scale parameter; and ***b*** was the growth-rate coefficient. Increases in ***a*** and decreases in ***b*** delay both the age at maximum growth rate (***a***/***b***) and the time required to reach 95% of ***M*** ((***a***+2.973)/***b***), resulting in a flatter growth curve. Conversely, lower ***a*** and higher ***b*** indicate faster growth. To evaluate model performance, we calculated the coefficient of determination (***R^2^***) using observed and predicted body weights, with values closer to 1 indicating better fit. Only individuals with ***R^2^*** > 0.90 were included in the heritability analyses of growth curve parameters.

### Estimating traits’ heritability and breeding values

The heritability of the body weight and the three growth curve parameters (***M***, ***a*** and ***b***) derived above were analyzed using a Monte Carlo-Markov Chain (MCMC) method implemented with the package **MCMCglmm** (Hadfield 2010) in R 4.5.1 (R Core Team 2022). All four traits were modeled as Gaussian-distributed variables. In general, ‘sex’ was included as a fixed effect to account for potential sex-specific effect, and ‘born_year’ was included as a random effect to capture variation associated with annual differences in breeding conditions (e.g. facility improvements, staff variation). Specifically, for body weight, the fixed effects included ‘day’ (age in days), ‘sex’ and the interaction between ‘day’ and ‘sex’ to account for possible sex-specific differences in growth. The MCMC chain was run for 100,000 iterations, with the first 20,000 iterations discarded as burn-in to eliminate potentially unstable estimates. A thinning interval of 100 was applied, retaining one draw every 100 iterations, resulting in a posterior sample of 800 draws for subsequent analyses. As for growth curve parameters (***M, a*** and ***b***), the fixed effect included ‘sex’ and the random effects were specified for ‘animal’ and ‘born_year’. Each MCMC chain was run for 1000,000 iterations, with a burn-in of 200,000 iterations and a thinning interval of 1,000. All models incorporated the population pedigree. Heritability (*h^2^*) was computed as the ratio of additive genetic variance to the total phenotypic variance, that is the ratio of the posterior variance component attributed to ‘animal’to the total phenotypic variance, where the latter included variance components of ‘animal’, ‘born_year’, and ‘residual’. Estimated breeding values of individual for each trait were obtained by extracting the posterior means of the ‘animal’random effects in each posterior sample from the fitted model.

The generational changes of breeding value were estimated following the approach described by Hadfield et al. (2010) to account for uncertainty and avoid anticonservative estimate of evolutionary change. Specifically, for each draw in the posterior sample we derived above, we calculated the mean breeding value of each generation by averaging the animal random effects. Then we regressed the generational mean estimated breeding values against generation to obtain the slope, so that each posterior sample yielded one estimate of the slope representing the temporal trend of estimated breeding values across generations. By synthesizing all posterior samples, we got the posterior distribution of the slope describing generational changes in estimated breeding values. We then computed the generational mean estimated breeding values within each posterior draw and synthesized them across all samples, enabling visualization of the overall temporal trend.

To assess whether the observed trend differed from expectations under genetic drift alone (i.e., in the absence of directional selection or evolution), we simulated data assuming no directional evolution for each posterior sample based on its additive genetic variance and pedigree structure. Then we calculated the generational mean estimated breeding values using the same manner and derived the posterior distribution of slope values expected under drift. By comparing the slope distributions from the observed and simulated data, we evaluated whether the observed estimated breeding values exhibited generational changes beyond what would be expected by drift alone. The proportion of posterior draws in which the observed slope exceeded the corresponding simulated slope was interpreted as a posterior predictive p-value. This value represents the probability that the observed trend in estimated breeding values cannot be explained solely by random genetic drift. Posterior predictive p-value close to 0 or 1 indicate that the observed trend is unlikely to result from drift alone and may reflect a genuine evolutionary response, whereas values near 0.5 suggest consistency with drift. To further assess whether the observed trend differed significantly from expectation, we calculated, for each posterior sample, the difference between the observed slope and the corresponding simulated slope. We then derived the 95% credible interval (CI) of this difference distribution. A trend was considered statistically significant when the 95% CI did not include zero.

### Predicting and monitoring individual health through offspring-parent growth trend comparisons

To monitor and predict offspring health, we compared the growth trend between the offspring and their parents. Assuming that both the body weight and growth trends are heritable, the mean parental growth trend can be used to predict the expected growth pattern of their offspring. Under normal and healthy development, an offspring’s daily growth trend should closely follow this parental mean. Deviations from the expected trajectory indicate potential health problems during the nestling period. Such deviations make unhealthy individuals readily identifiable. In captive breeding programs, early detection of these individuals allows timely veterinary intervention, improving their health and increasing the overall breeding success of the population.

We first calculated the correlation of body weight between mid-parents and the two groups of offspring (i.e., healthy vs. unhealthy). Parental body weight (***mid-parent***) was calculated by averaging the weights of both parents on the same day. If only one parent’s weight were available on a given day, the ***mid-parent*** value for that day would be recorded as *NA*. To ensure proper data for correlation analysis, individuals with fewer than five matching weight records with their parents were excluded. We ended up with a sub-dataset including 58 individuals, of which 47 were healthy and 10 were unhealthy. Paired Pearson correlation coefficients between offspring and parental weights during the development period were then calculated for different health groups.

Next, we employed the loess method within the **ggplot2** package (Wickham 2016) to visualize growth trends across groups using empirical data. These groups included the average weight of the parents (***mid-parent***, n = 17), the healthy group (***healthy***, n = 243) and the unhealthy group (***unhealthy***, n = 21). Again, assuming that body weight and daily growth trend are heritable and closely linked to body condition and nutritional status, we expect unhealthy individuals to deviate from the predicted growth pattern at specific time points. Such deviations would be observable in the plotted trajectories as departures from the parental-based expectations.

## Results

### General summary

The whole pedigree comprised 384 captive-bred individuals with known kin relationships from 2008 to 2025. Among them, 264 had more than 5 records of daily body weight during Day 0 to Day 42 of development, resulting in a total of 6422 daily weight records. Of these individuals, 102 were males, 97 were females and 65 remained unknown in gender. In terms of health status, 243 were classified as healthy and 21 as unhealthy. Gompertz growth curves were fitted for individuals with sufficient data. The maximum ***R^2^*** of the fitted curves was 0.9996, with an average of 0.9739, indicating generally high model performance. A total of 253 individuals achieved an ***R^2^*** greater than 0.90 (including 94 females and 96 males; 233 healthy ones and 20 unhealthy ones), which were subsequently used to estimate the heritability of the growth curve parameters.

### Heritability of body weight and growth curve parameters

Posterior estimates of variance components and heritability for body weight and three growth curve parameters were summarized in Table 1. The body weight exhibited significant heritability (*h^2^* = 0.195, 95% HPD: 0.139 ∼ 0.250). The main effect of ‘sex’ (posterior mean difference = −40.409, 95% CI: −56.874 ∼ −23.222, pMCMC < 0.001) and the day-by-sex interaction (posterior mean difference = 4.310, 95% CI: 3.704 ∼ 4.834, pMCMC < 0.001) were statistically significant, indicating that males exhibited a lower initial body weight and a relatively faster growth rates than females. For potential maximum weight ***M***, both the animal and birth-year components explained only a small proportion of the total variance, resulting in low heritability (*h^2^* = 0.023, 95% HPD: 0.00 ∼ 0.16). For growth time-scale parameter ***a*** (*h^2^*= 0.33, 95% HPD: 0.13 ∼ 0.53) and growth-rate coefficient ***b*** (*h^2^* = 0.21, 95% HPD: 0.10 ∼ 0.34), the results demonstrated significant heritability. Regarding sex differences, males had a significantly higher potential maximum weight ***M*** than females (posterior mean difference = 126.53, 95% CI: 60.29 ∼ 189.32, pMCMC < 0.001), while no significant sex effects were detected for growth time-scale parameter ***a*** or growth-rate coefficient ***b*** (pMCMC > 0.1).

**Table 1.**
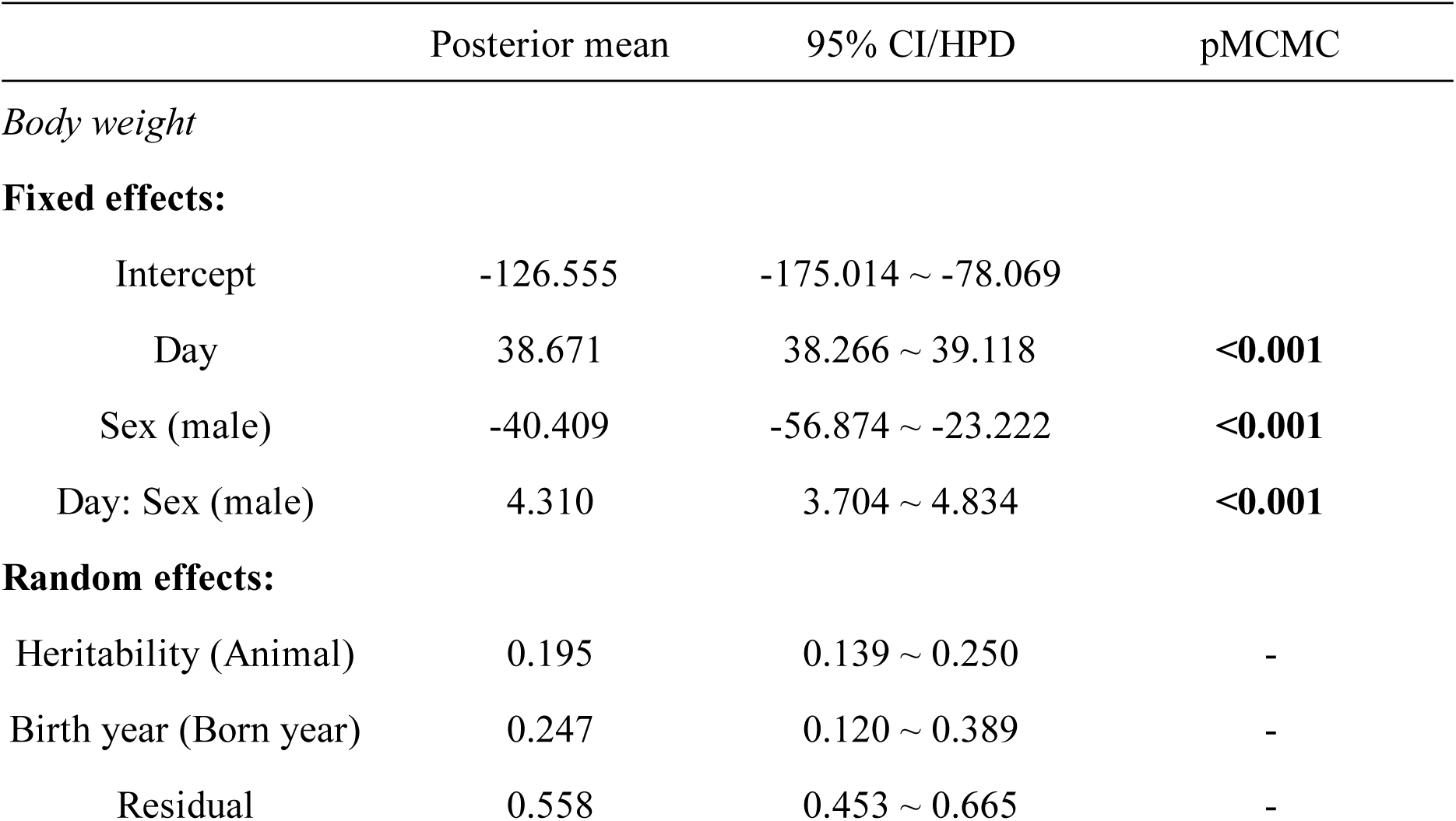
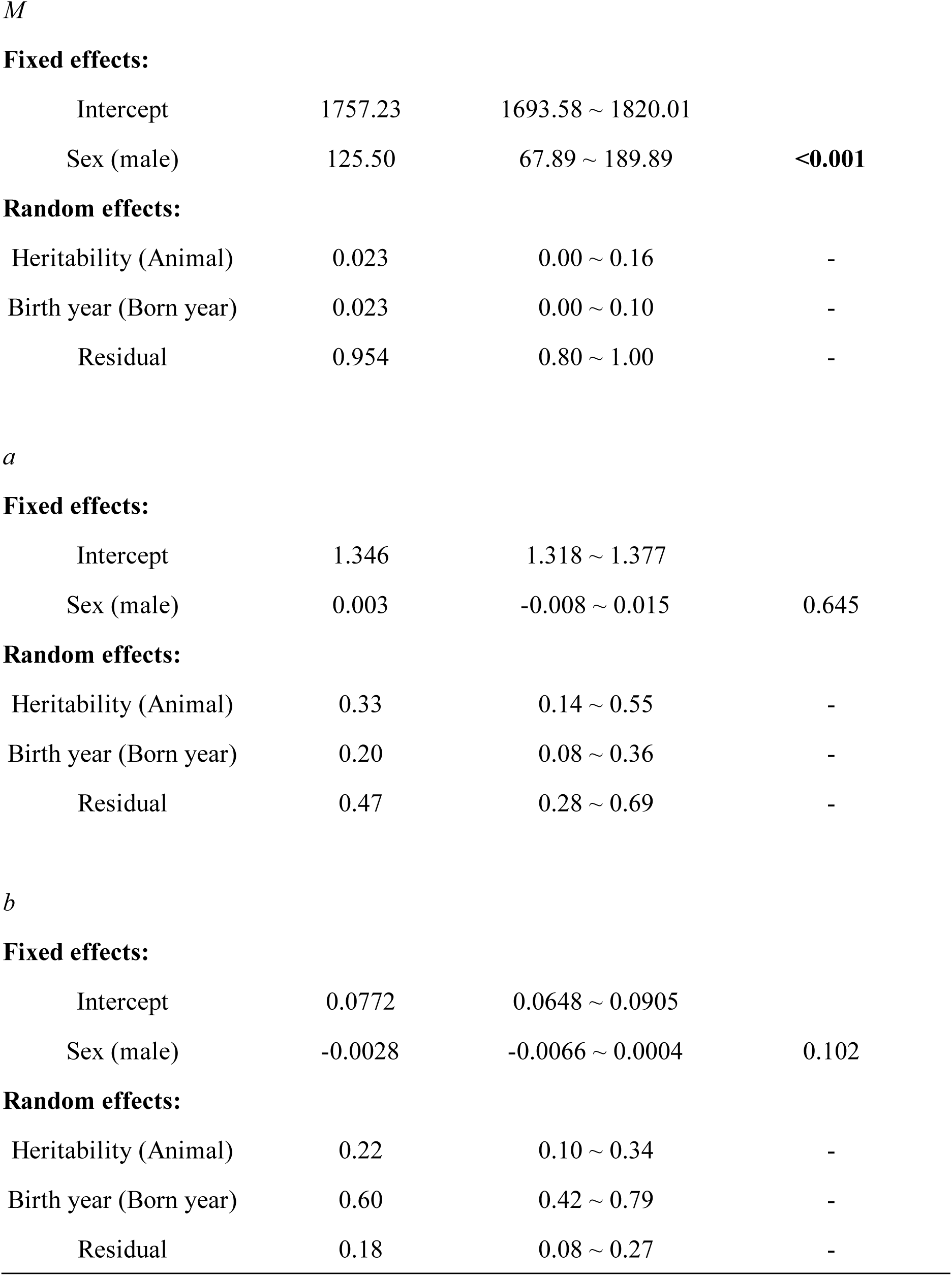
Posterior estimates of fixed effects with 95% credible interval (CI), and posterior mean value of heritability with 95% highest posterior density (HPD) intervals for body weight and growth curve parameters: ***M*** (potential maximum weight), ***a*** (growth time-scale parameter), ***b*** (growth-rate coefficient).

|  | Posterior mean | 95% CI/HPD | pMCMC |
| --- | --- | --- | --- |
| *Body weight* |  |  |  |
| **Fixed effects:** |  |  |  |
| Intercept | -126.555 | -175.014 ~ -78.069 |  |
| Day | 38.671 | 38.266 ~ 39.118 | <0.001 |
| Sex (male) | -40.409 | -56.874 ~ -23.222 | <0.001 |
| Day: Sex (male) | 4.310 | 3.704 ~ 4.834 | <0.001 |
| **Random effects:** |  |  |  |
| Heritability (Animal) | 0.195 | 0.139 ~ 0.250 | - |
| Birth year (Born year) | 0.247 | 0.120 ~ 0.389 | - |
| Residual | 0.558 | 0.453 ~ 0.665 | - |
| *M* |  |  |  |
| **Fixed effects:** |  |  |  |
| Intercept | 1757.23 | 1693.58 ~ 1820.01 |  |
| Sex (male) | 125.50 | 67.89 ~ 189.89 | <0.001 |
| **Random effects:** |  |  |  |
| Heritability (Animal) | 0.023 | 0.00 ~ 0.16 | - |
| Birth year (Born year) | 0.023 | 0.00 ~ 0.10 | - |
| Residual | 0.954 | 0.80 ~ 1.00 | - |
| *a* |  |  |  |
| **Fixed effects:** |  |  |  |
| Intercept | 1.346 | 1.318 ~ 1.377 |  |
| Sex (male) | 0.003 | -0.008 ~ 0.015 | 0.645 |
| **Random effects:** |  |  |  |
| Heritability (Animal) | 0.33 | 0.14 ~ 0.55 | - |
| Birth year (Born year) | 0.20 | 0.08 ~ 0.36 | - |
| Residual | 0.47 | 0.28 ~ 0.69 | - |
| *b* |  |  |  |
| **Fixed effects:** |  |  |  |
| Intercept | 0.0772 | 0.0648 ~ 0.0905 |  |
| Sex (male) | -0.0028 | -0.0066 ~ 0.0004 | 0.102 |
| **Random effects:** |  |  |  |
| Heritability (Animal) | 0.22 | 0.10 ~ 0.34 | - |
| Birth year (Born year) | 0.60 | 0.42 ~ 0.79 | - |
| Residual | 0.18 | 0.08 ~ 0.27 | - |

### Trait estimated breeding values of each individual and changes over generations

We examined the distributions of estimated breeding values for body-weight and growth curve parameters (Figure 2). For growth curve parameters, due to the low heritability of potential maximum weight ***M***, estimated breeding values were calculated only for the growth time-scale parameter ***a*** and growth-rate coefficient ***b***. Given the established association between larger body size and higher fitness, individuals with higher body weight and faster growth rates could be considered as ‘higher-quality’ individuals. Accordingly, estimated breeding values for body weight and the growth-rate coefficient ***b*** were ranked from highest to lowest, whereas values for the growth time-scale parameter ***a*** were ranked from lowest to highest. Vertical solid lines indicated the mean value for each trait, whereas dashed lines marked the top and bottom 5% of individuals, highlighting the extremes available for selective breeding and conservation of genetic resources. Full estimated breeding values for all individuals were provided in Table S1.

**Figure 2.**
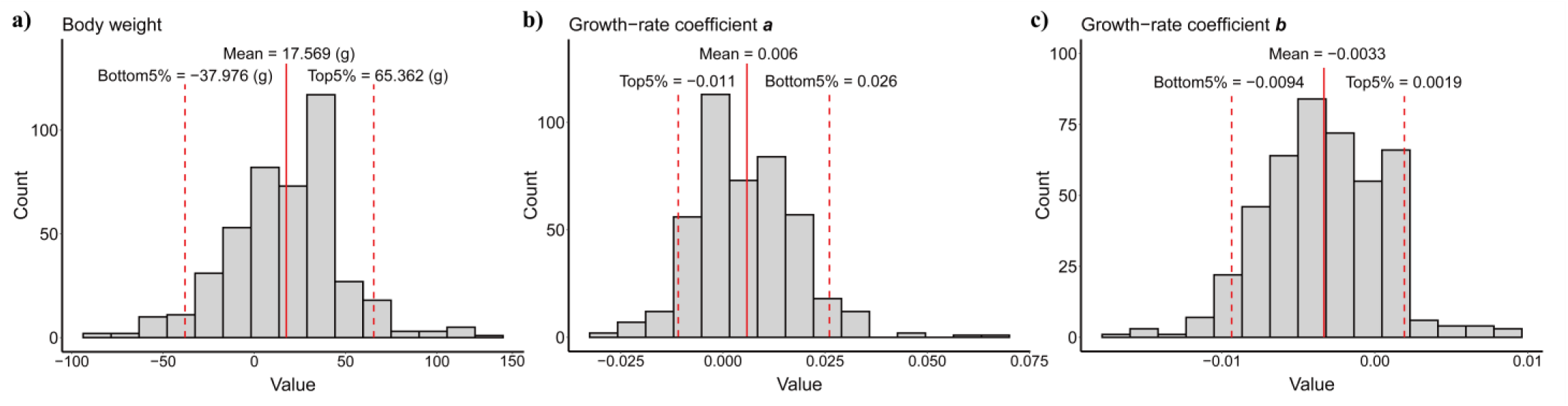
Distributions of estimated breeding values for a) body weight, b) growth time-scale parameter ***a*** and c) growth-rate coefficient ***b***. Vertical solid lines indicated the mean value for each trait, whereas dashed lines marked the top and bottom 5% values. Estimated breeding values were reported in the same units as the trait, reflecting the expected genetic deviation from the population mean. Negative values indicated a genetic contribution below the mean, whereas positive values indicated a contribution above the mean.

Examination of generational trends in posterior mean estimated breeding values showed that body weight exhibited a generally increasing pattern. As for the growth curve parameters, estimated breeding values for the growth time-scale parameter ***a*** showed a slight increasing trend with fluctuations, whereas those for the growth-rate coefficient ***b*** fluctuated but tended to decline across generations (Figure S2). Nevertheless, the posterior predictive p-values and 95% credible intervals indicated that none of them differed significantly from expectations under genetic drift (Body weight: posterior predictive p-value: 0.7625, 95% CI: −6.06 ∼ 13.53; growth time-scale parameter ***a***: posterior predictive p-value: 0.664, 95% CI: −0.004 ∼ 0.006; growth-rate coefficient ***b***: posterior predictive p-value: 0.376, 95% CI: −0.002 ∼ 0.002).

### Health predicting and monitoring through offspring-parent comparisons

Under the assumption that healthy offspring should closely follow the mean parental growth trend, deviations from this pattern can be used to identify individuals with poor health status (Figure 3a). Once such deviations are detected, timely veterinary interventions can be implemented.

**Figure 3.**
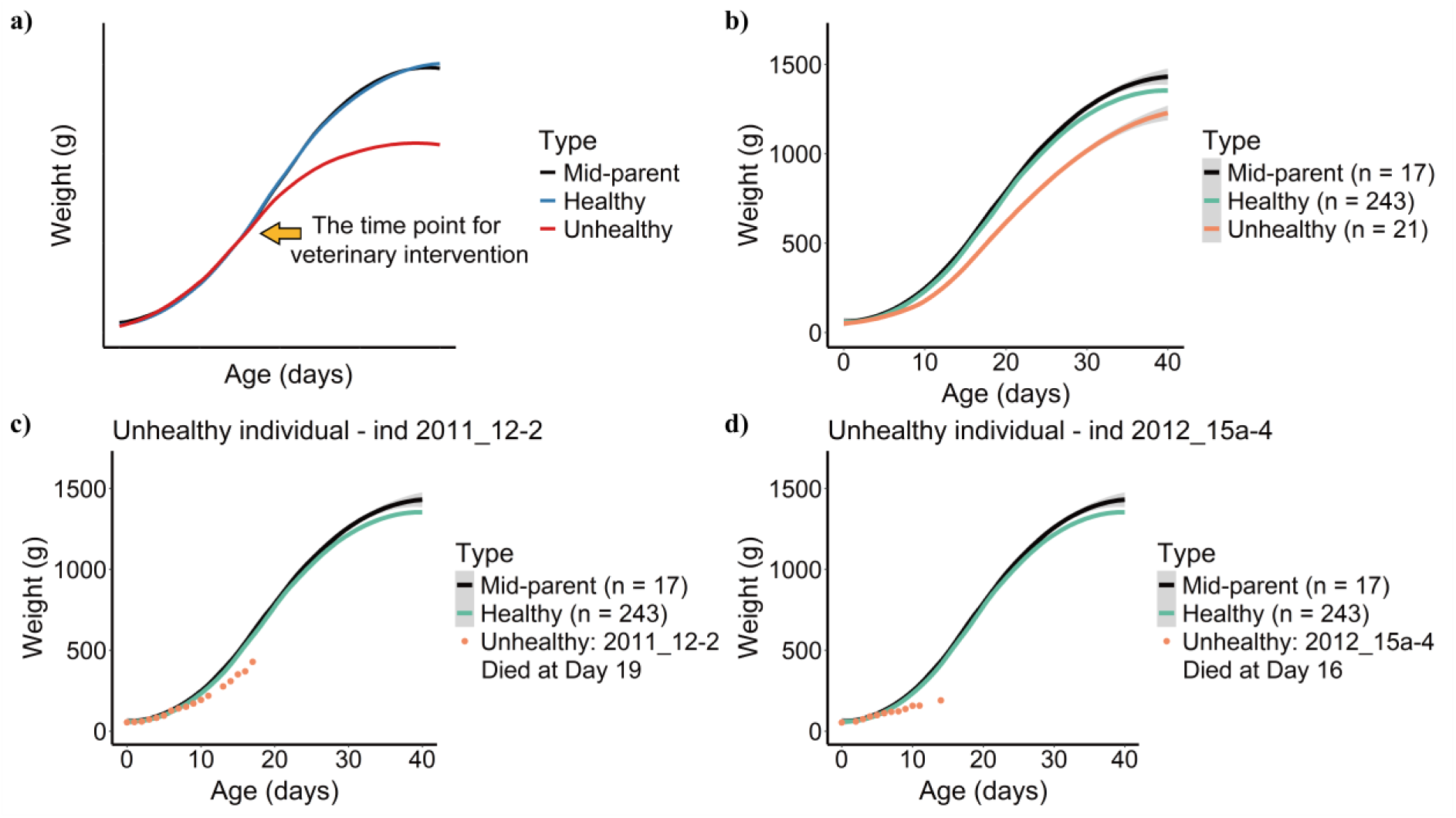
**a)** Schematic diagram of predicting and monitoring the offspring’s health based on parental performance to determine whether the offspring is normal and need veterinary intervention. Here, healthy offspring should behave similarly to their parents while the performance of unhealthy ones will deviate. The deviation time point is the moment when veterinary intervention is required; b) growth trend of individuals with different health statuses compared with their parents, based on empirical data in this study. Black curve: the mean growth trend of parents, green curve: the growth trend of healthy offspring, orange curve: the growth trend of unhealthy offspring, and pink curve: the growth trend of dead offspring; c) the unhealthy individual (2011_12-2), whose body weight was below normal starting around Day 9, ultimately died at Day 19; d) the unhealthy individual (2012_15a-4), whose body weight was below normal starting around Day 7, ultimately died at Day 16. The growth trend of all unhealthy individuals compared with parental performance and healthy individuals were shown in Figure S3.

Although Peason correlation coefficients did not differ significantly between healthy (0.975 ± 0.074, mean ± SD) and unhealthy individuals (0.965 ± 0.046) (*p* = 0.69), this lack of significance is likely attributable to the small sample size. Importantly, both individual-level patterns and group-level trends clearly demonstrated meaningful differences in growth performance. As shown in Figure 3b, healthy individuals exhibited growth trajectories that closely resemble those of their parents, whereas unhealthy individuals displayed consistently lower and visibly divergent growth curves. This deviation becomes even more pronounced in individuals that died early (Figure 3c ∼ 3d). Thus, despite the absence of statistically significant differences in correlation coefficients, the empirical growth trends provided a practical basis for early detection of unhealthy individuals and enabled timely health interventions.

## Discussion

In this study, we found that body weight and most growth curve parameters, except for potential maximum weight, of the Crested ibis during nestling period exhibited significant heritability (Table 1). Full estimated breeding values of all heritable traits were obtained for each individual (Table S1), providing a valuable reference for future conservation programs. Examination of estimated breeding values across eight generations indicated that the population appeared to have a higher body weight and grow slower, yet the observed trends did not differ significantly from expectations under genetic drift (Figure S2). Using parental performance to predict and monitor offspring health, we found that normally developing offspring showed growth curves closely resembling those of their parents, whereas unhealthy individuals, especially those who died early, displayed deviations from the expected developmental pattern (Figure 3). These results provide a practical framework for predicting individual growth and assessing health conditions during captive breeding. Early detection of abnormal growth allows timely veterinary intervention, thereby improving breeding outcomes and increasing management efficiency. The approaches demonstrated here can be extended to other species for which pedigrees and long-term phenotypic records are available.

In our study, we found significant heritability for body weight, growth time-scale parameter ***a*** and growth-rate coefficient ***b*** during the nestling development of the Crested ibis (Table 1), which was consistent with those reported across a range of bird species. For example, heritability estimates for body weight in Turkey (*Meleagris gallopavo*) averaged around 0.38 at 40, 60, 80 and 120 days (Aslam et al. 2011); in Great Lakes Piping plovers (*Charadrius melodus*), chick mass showed a heritability of 0.27 (Saunders and Cuthbert 2014) and in the Great tit (*Parus major*), fledging mass had a heritability of 0.29 (Garant et al. 2004). At the same time, previously studies of Partridges (*Rhynchotus rufescens*) showed that the heritability of asymptotic weight and rate of maturing were 0.22 and 0.12, respectively (Tholon et al. 2006), while estimates for growth curve parameters in meat-type chickens ranged from 0.25 to 0.48 (Mignon-Grasteau 1999). In contrast, the potential maximum weight ***M*** showed low heritability and a large residual variance in our analysis. This is likely attributable to limitations in the available dataset. Our weight records extended only to approximately 42 days of age, when the chicks typically fledge. Mass gain begins to slow but has not yet reached the final plateau. Individuals continue to gain weight from fledging until sexual maturity at around two years of age. For instance, in our records, females weighed on average 1248.5 g at 35 ∼ 42 days of age (n = 14) and males 1345 g (n = 13). According to ring-identification data from our captive population, post-fledging to one-year-old individuals averaged 1427 g for females (n = 11) and 1603 g for males (n = 14). For birds aged 1 ∼ 2 years, the averages were 1507 g for females (n = 11) and 1727 g for males (n = 15), and among individuals older than two years, females averaged 1482 g (n = 29) and males 1681 g (n = 37). In addition, weight records for some individuals were sparse. Together, the lack of data capturing the later weight plateau period likely introduced bias when estimating the potential maximum weight ***M*** from growth-curve models, result in a higher residual variance (Austin et al. 2011; Myhrvold 2013). In comparison, growth time-scale parameter ***a*** and growth-rate coefficient ***b*** are mainly determined by early growth trajectories and are therefore less affected by this limitation.

Apart from genetic factors, our results showed that sex has a measurable influence on body weight, while no significant sex differences were detected for the growth time-scale parameter ***a*** and growth-rate coefficient ***b***. This indicates that males and females may differ in absolute weight but follow similar growth trajectories during the nestling period. It is consistent with the fact that the Crested ibis is a sexually monomorphic and monogamous species with relatively weak sexual selection (Andersson 1994; Matheu et al. 2020). In addition, birth year had a pronounced effect on body weight, which is likely linked to interannual variation in maternal nutritional condition. Maternal condition is known to influence egg size, which in turn affects chick birth weight and early developmental performance (Bolton et al. 1992; Williams 1994; Grindstaff et al. 2005; Krist 2011). Birth year also contributed to variation in the growth time-scale parameter ***a*** and growth-rate coefficient ***b***. Growth time-scale parameter ***a*** primarily determines the timing of transition into the rapid growth phase, whereas growth - rate coefficient ***b*** mainly governs the growth rate during this phase. Given that feeding conditions and the completeness of facilities likely varied among years, these effects are consistent with evidence that nutritional conditions during the nestling period can influence growth rates (Negro et al. 1994).

In our work, the estimated breeding values of for body weight and growth curve parameters (the growth time-scale parameter ***a*** and growth-rate coefficient ***b***) remained generally consistent across generations. The posterior distribution of the intergenerational slope did not differ significantly from those generated under drift-only simulations (Figure S2), suggesting that clear directional evolutionary changes have not yet occurred. This stability is encouraging for the Dongzhai population, as it suggests that long-term captive breeding has not markedly altered growth-related traits. Still, some caution is needed because sample sizes vary among cohorts and most individuals belong to early generations, which may limit the detection of subtle changes. Additionally, environmental variation may also mask phenotypic responses (Merilä et al. 2001). As more generations accumulate, clearer trends may emerge. Despite the overall stability, body weight and growth time-scale parameter ***a*** showed a slight increase, whereas growth-rate coefficient ***b*** tended to decrease (Figure S2). Together, these shifts would produce a higher body weight with a flatter growth curve, implying a slower overall growth rate during the nestling period. This pattern is consistent with the low-stress conditions of captivity compared with the selective pressures of the wild (Remes and Martin 2002; Merrill et al. 2021). Overall, the lack of marked generational change suggests that current captive breeding practices have largely preserved natural growth profiles, which is important for maintaining the species’ reintroduction potential.

Since body weight, growth time-scale parameter ***a*** and growth-rate coefficient ***b*** showed significant heritability in Crested ibis, these traits can be used to predict offspring performances. We’ve assumed that the nestling growth trend of an offspring can be predicted by the mean of its parents’ growth trend, and the performances of healthy ones should be similar to that of their parents, while those who are unhealthy can be detected when their growth curves deviated and have necessary interventions applied (Figure 3a). Our results showed that the weight of healthy individuals had a relatively higher correlation with their parents than that of unhealthy individuals, although the difference was not statistically significant, likely due to limited overlapping measurements among parents and offspring. And in some unhealthy individuals, only the period before deviation had corresponding records, which may have obscured differences in correlation estimates. Nevertheless, the growth curves themselves revealed clear deviations. At both the individual and group levels, unhealthy nestlings displayed growth trends that diverged from parental and healthy-offspring patterns, particularly among those who died early (Figure 3b-3d & Figure S3). Given the long breeding years of this species (usually sexually mature in the third year and breed every year with a long-lived lifespan) (Yu et al. 2007), which means the same parent can produce multiple times, our findings provide a practical way for predicting normal growth expectations and identifying abnormal individuals early in captivity, thereby improving the conservation efficiency of this endangered species. Similar work has been done on another endangered avian species, kākāpō ( *Strigops habroptilus*). Researches based on long-term pedigree and phenotypic records revealed that declines in chick growth curves preceded clinical signs of aspergillosis, enabling intervention at the onset of deviation (Guhlin et al. 2023). Overall, with long-term pedigree and phenotypic datasets, this framework can be readily applied to other endangered species and may contribute to efforts aimed at slowing biodiversity loss.

Our analyses were based on phenotypic data with the population pedigree, and it is necessary to integrate them with more genetic and genomic data in future studies (Wright et al. 2021). As an endangered species restored from a small founder population of only 7 individuals, with just 2 breeding pairs, the Crested ibis has experienced persistent inbreeding. Given that inbreeding is known to cause outcomes such as embryonic mortality, low genetic diversity, and promote the accumulation of deleterious alleles (Feng et al. 2019; Fu et al. 2019), it is possible that the unhealthy individuals detected in our study were similarly affected by inbreeding. Such problems cannot be fully resolved through routine veterinary interventions alone, and the true severity of inbreeding may be underestimated because founder relatedness remains unknown in the absence of detailed genetic data. Incorporating genomic information would provide more accurate estimates of relatedness, thereby improving quantitative genetic analyses (Henkel et al. 2012; Bergner et al. 2014). Genomic tools also enable the identification of loci associated with key traits, offering deeper insight into the genetic basis of phenotypic variation (Guhlin et al. 2023; Strickland et al. 2024). Although genomic research on the Crested ibis has progressed steadily (Zhang et al. 2004; Li et al. 2014; Fu et al. 2019; Choi et al. 2020), large-scale integration of genomic and long-term phenotypic datasets is still lacking. Future efforts that combine these complementary sources of information will improve strategies for minimizing inbreeding, clarify the mechanisms governing population recovery, and enhance our ability to address key conservation challenges. Such advances will ultimately support the sustainable recovery of the Crested ibis, and facilitate the true return of these animals into the wild.

## Conclusion

In this study, we showed that the body weight and key growth curve parameters in the Crested ibis during nestling development exhibit significant heritability, suggesting their utility for predicting offspring performance and informing future breeding management. The body weight showed an overall increasing trend accompanied by slower growth rates across eight generations, though the observed changes did not differ significantly from drift expectations, which aligns with conservation goals that aim to maintain the species’ natural characteristics. By comparing individuals with different health statuses, we further show that unhealthy offspring, particularly those who died early, display atypical growth trajectories characterized by abnormal inflection points that deviate from parental expectations, suggesting that such deviations can serve as indicators for targeted veterinary care. Overall, our work confirms that body weight and key growth curve parameters of Crested ibis are heritable and can be used to monitor the health status of Crested ibis and guide future breeding, thereby enhancing the conservation efficiency. This practical framework is also readily applicable to other endangered species wherever long-term pedigree and phenotypic datasets are available.

## References

Abou Khadiga G, Mahmoud BYF, El-Full EA (2016) Genetic evaluation of early egg production and maturation traits using two different approaches in Japanese quail. Poult Sci 95:774–779. 10.3382/ps/pev386

Anderson A (1984) Irreproducing ibis. Nature 309:741–741. 10.1038/309741b0

Andersson M (1994) Sexual Selection. Princeton University Press

Archibald GW, Lantis SDH (1979) Conservation of the Japanese Crested Ibis. Proceedings of the Colonial Waterbird Group 2:1. 10.2307/1520927

Archibald GW, Lantis SDH, Lantis LR, Munetchika I (1980) Endangered ibises *Threskiornithinae:* their future in the wild and in captivity. International Zoo Yearbook 20:6–17. 10.1111/j.1748-1090.1980.tb00936.x

Aslam ML, Bastiaansen JW, Crooijmans RP, et al (2011) Genetic variances, heritabilities and maternal effects on body weight, breast meat yield, meat quality traits and the shape of the growth curve in turkey birds. BMC Genet 12:14. 10.1186/1471-2156-12-14

Austin SH, Robinson TR, Robinson WD, Ricklefs RE (2011) Potential biases in estimating the rate parameter of sigmoid growth functions. Methods Ecol Evol 2:43–51. 10.1111/j.2041-210X.2010.00055.x

Ballou JD, Foose TJ, Lacy RC, Seal US (1989) Florida Panther Felis concolor coryi Population Viability Analysis and Recommendations. Captive breeding Specialist Group Species Survival Commission IUCN

Beissinger SR, Wunderle JM, Meyers JM, et al (2008) Anatomy of a Bottleneck: Diagnosing Factors Limiting Population Growth in the Puerto Rican Parrot. Ecol Monogr 78:185– 203. 10.1890/07-0018.1

Bergner LM, Jamieson IG, Robertson BC (2014) Combining genetic data to identify relatedness among founders in a genetically depauperate parrot, the Kakapo ( *Strigops habroptilus*). Conserv Genet 15:1013–1020. 10.1007/s10592-014-0595-y

BirdLife-International (2001) Threatened Birds of Asia: the BirdLife International Red Data Book. BirdLife International, Cambridge., Cambridge

Boag PT (1987) Effects of Nestling Diet on Growth and Adult Size of Zebra Finches (Poephila guttata). The Auk 104:155–166. 10.1093/auk/104.2.155

Bolton M, Houston D, Monaghan P (1992) Nutritional Constraints on Egg Formation in the Lesser Black-Backed Gull: An Experimental Study. J Anim Ecol 61:521–532. 10.2307/5607

Cai D, Huang H, Huang Z, et al (2024) Population and Distribution Status of Reintroduced Crested Ibis in Dongzhai Reserve, Henan Province (in Chinese). Chinese Journal of Wildlife 45:409–414

China National Radio (2025) The world’s oldest crested ibis has reached the age of 39! (in Chinese). https://www.cnr.cn/erwen/ewbfycsy/?id=64&lastNewsId=29854259. Accessed 26 Apr 2025

Choi EH, Kim G, Baek SY, et al (2020) Development and Characterization, and Application of Ten Polymorphic Microsatellite Markers in the Crested Ibis *Nipponia nippon* from South Korea. Animal Systematics, Evolution and Diversity 36:154–158. 10.5635/ASED.2020.36.2.029

Cowie RH, Bouchet P, Fontaine B (2022) The Sixth Mass Extinction: fact, fiction or speculation? Biol Rev 97:640–663. 10.1111/brv.12816

Ding C (2004) Research on the Crested Ibis (in Chinese). Shanghai Scientific and Technological Educational Publishing House, Shanghai, China

Ding C, Ma Z, Li X, et al (1999) A preliminary study on the movements of the young crested ibis (Nipponia nippon). In: Proceedings of the International Crested Ibis Conservation Symposium. China Forestry Publishing House, Shaanxi, China

Donnelly RE, Sullivan KA (1998) Foraging Proficiency and Body Condition of Juvenile American Dippers. The Condor 100:385–388. 10.2307/1370282

Feng S, Fang Q, Barnett R, et al (2019) The Genomic Footprints of the Fall and Recovery of the Crested Ibis. Curr Biol 29:340–349.e7. 10.1016/j.cub.2018.12.008

Fu CZ, Guang XM, Wan QH, Fang SG (2019) Genome Resequencing Reveals Congenital Causes of Embryo and Nestling Death in Crested Ibis (*Nipponia nippon*). Genome Biol Evol 11:2125–2135. 10.1093/gbe/evz149

Garant D, Kruuk LEB, McCleery RH, Sheldon BC (2004) Evolution in a Changing Environment: A Case Study with Great Tit Fledging Mass. The American Naturalist 164:E115–E129. 10.1086/424764

Grindstaff J, Demas GE, Ketterson ED (2005) Diet quality affects egg size and number but does not reduce maternal antibody transmission in Japanese quail *Coturnix japonica*. J Anim Ecol 74:1051–1058. 10.1111/j.1365-2656.2005.01002.x

Grueber CE, Laws RJ, Nakagawa S, Jamieson IG (2010) Inbreeding depression accumulation across life-history stages of the endangered Takahe. Conserv Biol 24:1617–1625. 10.1111/j.1523-1739.2010.01549.x

Guhlin J, Le Lec MF, Wold J, et al (2023) Species-wide genomics of kākāpō provides tools to accelerate recovery. Nat Ecol Evol 7:1693–1705. 10.1038/s41559-023-02165-y

Hadfield JD (2010) MCMC Methods for Multi-Response Generalized Linear Mixed Models: The MCMCglmm R Package. Journal of Statistical Software 33:1–22. 10.18637/jss.v033.i02

Hadfield JD, Wilson AJ, Garant D, et al (2010) The Misuse of BLUP in Ecology and Evolution. The American Naturalist 175:116–125. 10.1086/648604

Henkel JR, Jones KL, Hereford SG, et al (2012) Integrating microsatellite and pedigree analyses to facilitate the captive management of the endangered Mississippi sandhill crane ( *Grus canadensis pulla*). Zoo Biol 31:322–335. 10.1002/zoo.20399

Huang Z, Zhang J, Tang S, et al (2006) The growth of nestling of crested ibis hand-rearing (in Chinese). Chinese Journal of Zoology 87–92

Huang Z, Zhu J, Wang K (2016) The study on artificial propagation of crested ibis in Dongzhai Natural Protection Area of Henan province (in Chinese). Contemporary Animal Husbandry 29–32

Isbell F, Balvanera P, Mori AS, et al (2023) Expert perspectives on global biodiversity loss and its drivers and impacts on people. Front Ecol Environ 21:94–103. 10.1002/fee.2536

IUCN (2025) The IUCN Red List of Threatened Species. Version 2025–2. https://www.iucnredlist.org. Accessed on 14 October 2025. https://www.iucnredlist.org/en.

Khan A, Patel K, Shukla H, et al (2021) Genomic evidence for inbreeding depression and purging of deleterious genetic variation in Indian tigers. Proceedings of the National Academy of Sciences 118:e2023018118. 10.1073/pnas.2023018118

Killpack TL, Karasov WH (2012) Growth and development of house sparrows (*Passer domesticus*) in response to chronic food restriction throughout the nestling period. J Exp Biol 215:1806–1815. 10.1242/jeb.066316

Krauss J, Bommarco R, Guardiola M, et al (2010) Habitat fragmentation causes immediate and time-delayed biodiversity loss at different trophic levels. Ecology Letters 13:597–605. 10.1111/j.1461-0248.2010.01457.x

Krementz DG, Nichols JD, Hines JE (1989) Postfledging survival of European starlings. Ecology 70:646–655. 10.2307/1940216

Krist M (2011) Egg size and offspring quality: a meta-analysis in birds. Biol Rev Camb Philos Soc 86:692–716. 10.1111/j.1469-185X.2010.00166.x

Kruuk LEB (2004) Estimating genetic parameters in natural populations using the ‘animal model.’ Phil Trans R Soc Lond B 359:873–890. 10.1098/rstb.2003.1437

Lambeck RJ (1997) Focal Species: A Multi - Species Umbrella for Nature Conservation: Especies Focales: Una Sombrilla Multiespec í fica para Conservar la Naturaleza. Conserv Biol 11:849–856. 10.1046/j.1523-1739.1997.96319.x

Li F, Huang S (1986) Investigation on the reproductive habits of crested ibis (in Chinese). Bulletin of Biology 6–8

Li H (2023) Big increase: Crested ibis population exceeds 10,000 globally after 42 years of protection. https://greenchina.chinadaily.com.cn/s/202311/05/WS655b4c0d498ed2d7b7ea07f9/big-increase-crested-ibis-population-exceeds-10-000-globally-after-42-years-of-protection.html. Accessed 15 Oct 2025

Li S, Li B, Cheng C, et al (2014) Genomic signatures of near-extinction and rebirth of the crested ibis and other endangered bird species. Genome Biol 15:557. 10.1186/s13059-014-0557-1

Li X, Tian H, Li D (2009) Why the crested ibis declined in the middle twentieth century. Biodivers Conserv 18:2165–2172. 10.1007/s10531-009-9580-z

Liu Y (1981) Rediscovery of the crested ibis in Qin Mountain (in Chinese). Acta Zoologica Sin 27:273

Mallard BA, Wilkie BN, Kennedy BW, Quinton M (1992) Use of estimated breeding values in a selection index to breed Yorkshire pigs for high and low immune and innate resistance factors. Anim Biotechnol 3:257–280. 10.1080/10495399209525776

Malm S, Fikse WF, Danell B, Strandberg E (2008) Genetic variation and genetic trends in hip and elbow dysplasia in Swedish Rottweiler and Bernese Mountain Dog. J Anim Breed Genet 125:403–412. 10.1111/j.1439-0388.2008.00725.x

Manjula P, Park H-B, Seo D, et al (2018) Estimation of heritability and genetic correlation of body weight gain and growth curve parameters in Korean native chicken. Asian-australas J Anim Sci 31:26–31. 10.5713/ajas.17.0179

Matheu E, del Hoyo J, Kirwan GM, Garcia E (2020) Crested Ibis (*Nipponia nippon*), version 1.0. In: Hoyo J del, Elliott A, Sargatal J, et al. (eds) Birds of the World. Cornell Lab of Ornithology, Ithaca, NY, USA

Merilä J, Kruuk LE, Sheldon BC (2001) Cryptic evolution in a wild bird population. Nature 412:76–79. 10.1038/35083580

Merilä J, Svensson E (1997) Are Fat Reserves in Migratory Birds Affected by Condition in Early Life? J Avian Biol 28:279–286. 10.2307/3676940

Merrill L, Jones TM, Brawn JD, Ward MP (2021) Early-life patterns of growth are linked to levels of phenotypic trait covariance and postfledging mortality across avian species. Ecol Evol 11:15695–15707. 10.1002/ece3.8231

Mignon-Grasteau S (1999) Genetic parameters of growth curve parameters in male and female chickens. Br Poult Sci 40:44–51. 10.1080/00071669987827

Monaghan P, Uttley JD, Burns MD, et al (1989) The Relationship Between Food Supply, Reproductive Effort and Breeding Success in Arctic Terns Sterna paradisaea. The Journal of Animal Ecology 58:261. 10.2307/4999

Morrissey MB (2018) pedantics: Functions to Facilitate Power and Sensitivity Analyses for Genetic Studies of Natural Populations. R package version 1.7. https://CRAN.R-project.org/package=pedantics. Accessed 31 Oct 2025

Myhrvold NP (2013) Revisiting the Estimation of Dinosaur Growth Rates. PLOS ONE 8:e81917. 10.1371/journal.pone.0081917

Naef-Daenzer B, Grüebler MU (2008) Post-Fledging Range use of Great Tit *Parus major* Families in Relation to Chick Body Condition. Ardea 96:181–190. 10.5253/078.096.0204

Narinç D, Aksoy T, Kaplan S (2016) Effects of Multi-Trait Selection on Phenotypic and Genetic Changes in Japanese Quail (*Coturnix coturnix japonica*). Jpn Poult Sci 53:103– 110. 10.2141/jpsa.0150068

Narinc D, Karaman E, Aksoy T, Firat MZ (2014) Genetic parameter estimates of growth curve and reproduction traits in Japanese quail. Poult Sci 93:24–30. 10.3382/ps.2013-03508

Narinç D, Narinç NÖ, Aygün A (2017) Growth curve analyses in poultry science. World’s Poult Sci J 73:395–408. 10.1017/S0043933916001082

N’Dri AL, Mignon-Grasteau S, Sellier N, et al (2006) Genetic relationships between feed conversion ratio, growth curve and body composition in slow-growing chickens. Br Poult Sci 47:273–280. 10.1080/00071660600753664

Negro JJ, Chastin A, Bird DM (1994) Effects of Short-Term Food Deprivation on Growth of Hand-Reared American Kestrels. The Condor: Ornithological Applications 96:749–760. 10.2307/1369478

Okahisa Y, Kaneko Y, Nagata H, Ozaki K (2022) Effects of Rearing Methods on the Reproduction of Reintroduced Crested Ibis *Nipponia nippon* on Sado Island, Japan. Ornithological Science 21:. 10.2326/osj.21.145

Qiu G, Jiang J, Bai H, et al (2023) Reproductive Behavior of the Released Crested Ibis in Deqing, Zhejiang Province (in Chinese). Chinese Journal of Zoology 58:348–356

R Core Team (2022) R: A language and environment for statistical computing. R Foundation for Statistical Computing, Vienna, Austria

Regan CE, Tuke LA, Colpitts J, et al (2019) Evolutionary quantitative genetics of juvenile body size in a population of feral horses reveals sexually antagonistic selection. Evol Ecol 33:567–584. 10.1007/s10682-019-09988-x

Remes V, Martin TE (2002) Environmental influences on the evolution of growth and developmental rates in passerines. Evolution 56:2505–2518. 10.1111/j.0014-3820.2002.tb00175.x

Roberge J, Angelstam P (2004) Usefulness of the Umbrella Species Concept as a Conservation Tool. Conserv Biol 18:76–85. 10.1111/j.1523-1739.2004.00450.x

Ronget V, Gaillard J, Coulson T, et al (2018) Causes and consequences of variation in offspring body mass: meta - analyses in birds and mammals. Biol Rev 93:1 – 27. 10.1111/brv.12329

Saunders SP, Cuthbert FJ (2014) Genetic and environmental influences on fitness-related traits in an endangered shorebird population. Biol Conserv 177:26–34. 10.1016/j.biocon.2014.06.005

Seddon PJ, Griffiths CJ, Soorae PS, Armstrong DP (2014) Reversing defaunation: Restoring species in a changing world. Science 345:406–412. 10.1126/science.1251818

Shi D, Wang K, Yu X (1999) Status and suggestions of conservation and research of crested ibis (in Chinese). In: Proceedings of the International Crested Ibis Conservation Symposium. China Forestry Publishing House, Shaanxi, China

Shi D, Yu X (1989) The breeding habits of the crested ibis (*Nipponia nippon*) (in Chinese). Zoological Research 10:327–332

Singh MK, Kumar S, Sharma RK, et al (2018) Heritability Estimates of Adult Body Weight and Egg Production Traits in Indigenous Uttara Chickens. biop 10:357. 10.9735/0975-2862.10.2.357-359

Strickland K, Matthews B, Jónsson ZO, et al (2024) Microevolutionary change in wild stickleback: Using integrative time-series data to infer responses to selection. Proceedings of the National Academy of Sciences 121:e2410324121. 10.1073/pnas.2410324121

Tarwater CE, Ricklefs RE, Maddox JD, Brawn JD (2011) Pre-reproductive survival in a tropical bird and its implications for avian life histories. Ecology 92:1271 – 1281. 10.1890/10-1386.1

Tholon P, Freitas EC, Stein MS, Queiroz SA (2006) Genetic parameters estimates to Gompertz growth curve parameters fitted to partridges (Rhynchotus rufescens) raised in captivity. In: EPC 2006 - 12th European Poultry Conference. Verona, Italy, p 78

Toghiani S (2012) Quantitative Genetic Application in the Selection Process for Livestock Production. In: Javed K (ed) Livestock Production. InTech

Uchida Y (1970) On the color change in Japanese Crested Ibis. Journal of the Yamashina Institute for Ornithology 6:54–72. 10.3312/jyio1952.6.54

Urban MC (2015) Accelerating extinction risk from climate change. Science 348:571–573. 10.1126/science.aaa4984

Wajiki Y, Kaneko Y, Sugiyama T, et al (2014) Demographic Analyses in the Japanese Captive Population of Japanese Crested Ibis (*Nipponia nippon*). Jpn J Zoo Wildl Med 19:57– 67. 10.5686/jjzwm.19.57

Wang K, Shi D (1999) Observation of anniversary and daily activity of the crested ibis (in Chinese). In: Proceedings of the International Crested Ibis Conservation Symposium. China Forestry Publishing House, Shaanxi, China

Wendeln H, Becker PH (1999) Effects of parental quality and effort on the reproduction of common terns. J Anim Ecol 68:205–214. 10.1046/j.1365-2656.1999.00276.x

Wickham H (2016) ggplot2: Elegant Graphics for Data Analysis. Springer International Publishing

Williams TD (1994) Intraspecific variation in egg size and egg composition in birds: effects on offspring fitness. Biol Rev Camb Philos Soc 69:35–59. 10.1111/j.1469-185x.1994.tb01485.x

Wilson AJ, Pemberton JM, Pilkington JG, et al (2007) Quantitative genetics of growth and cryptic evolution of body size in an island population. Evol Ecol 21:337. 10.1007/s10682-006-9106-z

Wilson AJ, Réale D, Clements MN, et al (2010) An ecologist’s guide to the animal model. J Anim Ecol 79:13–26. 10.1111/j.1365-2656.2009.01639.x

Wingfield JC, Ishii S, Kikuchi M, et al (2000) Biology of a critically endangered species, the Toki (Japanese Crested Ibis) *Nipponia nippon*. Ibis 142:1–11. 10.1111/j.1474-919X.2000.tb07677.x

Winsor CP (1932) The Gompertz Curve as a Growth Curve. Proc Natl Acad Sci USA 18:1–8. 10.1073/pnas.18.1.1

Wright BR, Hogg CJ, McLennan EA, et al (2021) Assessing evolutionary processes over time in a conservation breeding program: a combined approach using molecular data, simulations and pedigree analysis. Biodivers Conserv 30:1011–1029. 10.1007/s10531-021-02128-4

Xie P, Zhao H, Liu D, et al (2020) Acclimation of Crested Ibis (*Nipponia nippon*) in Beidaihe wetland (in Chinese). Chinese Journal of Ecology 39:180–185

Yu X, Li X, Huo Z (2015) Breeding ecology and success of a reintroduced population of the endangered Crested Ibis *Nipponia nippon*. Bird Conserv Int 25:207–219. 10.1017/S0959270914000136

Yu X, Lu B, Lu X, Liu N (2007) Influences of age on the reproductive success of the crested ibis *Nipponia nippon* (in Chinese). Acta Zoologica Sinica 53:812–818

Yu X, Xi Y, Lu B, et al (2010) Postfledging and Natal Dispersal of Crested Ibis in the Qinling Mountains, China. The Wilson Journal of Ornithology 122:228–235

Zacharias MA, Roff JC (2001) Use of focal species in marine conservation and management: a review and critique. Aquat Conserv Mar Freshwater Ecosyst 11:59–76. 10.1002/aqc.429

Zhai T, Lu B, Zhang Y, et al (1999) Study of the reproductive ecology of crested ibis Nipponia nippon (in Chinese). In: Proceedings of the International Crested Ibis Conservation Symposium. China Forestry Publishing House, Shaanxi, China

Zhai T, Lu X, Lu B, et al (2011) Nest building, egglaying, hatching, and breeding of crested ibis ( *Nipponia nippon*) (in Chinese). Acta Zoologica Sinica 47:508–511

Zhang B, Fang S-G, Xi Y-M (2004) Low genetic diversity in the Endangered Crested Ibis *Nipponia nippon* and implications for conservation. Bird Conserv Int 14:183–190. 10.1017/S0959270904000231

Zheng J, Rees-Baylis E, Janzen T, et al (2024) Inbreeding and demography interact to impact population recovery from bottlenecks

Zheng L, Wang Y, Zhu J, et al (2018) Habitat Evaluation for Reintroduced Crested Ibis (*Nipponia nippon*) in Dongzhai National Nature Reserve, China, Based on a Maximum Entropy Model. Pakistan Journal of Zoology 50:. 10.17582/journal.pjz/2018.50.4.1319.1327

