## Supplementary material for "Monitor individual health and improve breeding success in Crested ibis (*Nipponia nippon*)": Figure S1

Figure S1. The pedigree of the captive population used in this study. Each black dot represents an individual. Blue lines radiating from a dot indicate that the corresponding individual participated in breeding as a male, while red lines indicate participation as a female. The endpoints of the lines represent the individual produced by them.

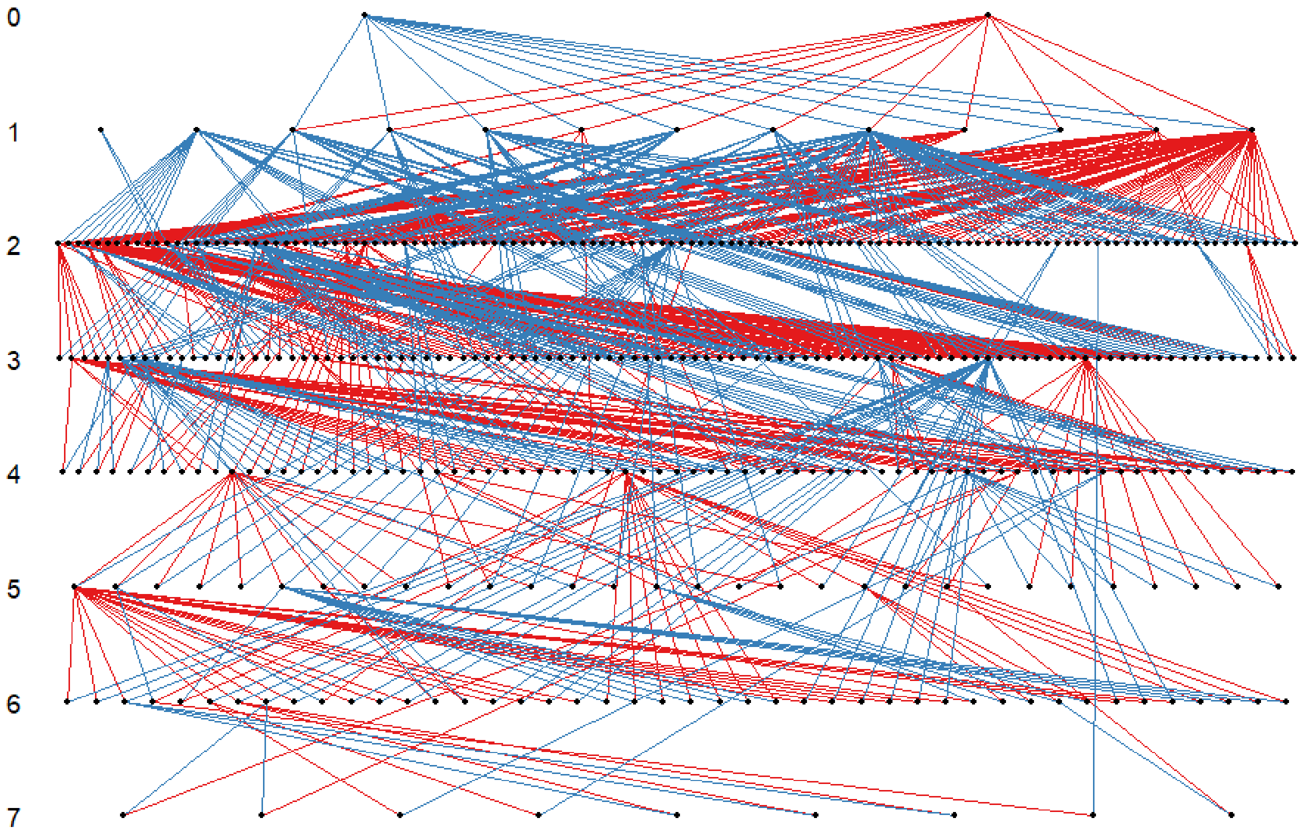
