## Supplementary material for "Monitor individual health and improve breeding success in Crested ibis (*Nipponia nippon*)": Figure S2

Figure S2. Generational trends of estimated breeding values and posterior slopes distributions. a) generational trend of estimated breeding values for body weight; b) the posterior slope distributions of estimated breeding values for body weight; c) generational trend of estimated breeding values for growth time-scale parameter ***a***; d) the posterior slope distributions of estimated breeding values for growth time-scale parameter ***a***; e) generational trend of estimated breeding values for growth-rate coefficient ***b***; f) the posterior slope distributions of estimated breeding values for growth-rate coefficient ***b***. In panels a), c) and e), the black solid line represents the mean breeding value across generations, with the grey shaded area indicating the 95% credible interval (CI). In panels b), d) and f), solid lines represent the slopes of estimated breeding values based on observed data, dashed lines represent those from simulated data under genetic drift, and red vertical lines indicate the mean of each distribution. A positive slope indicates an increasing trend in estimated breeding values across generations, whereas a negative slope indicates a decreasing trend.

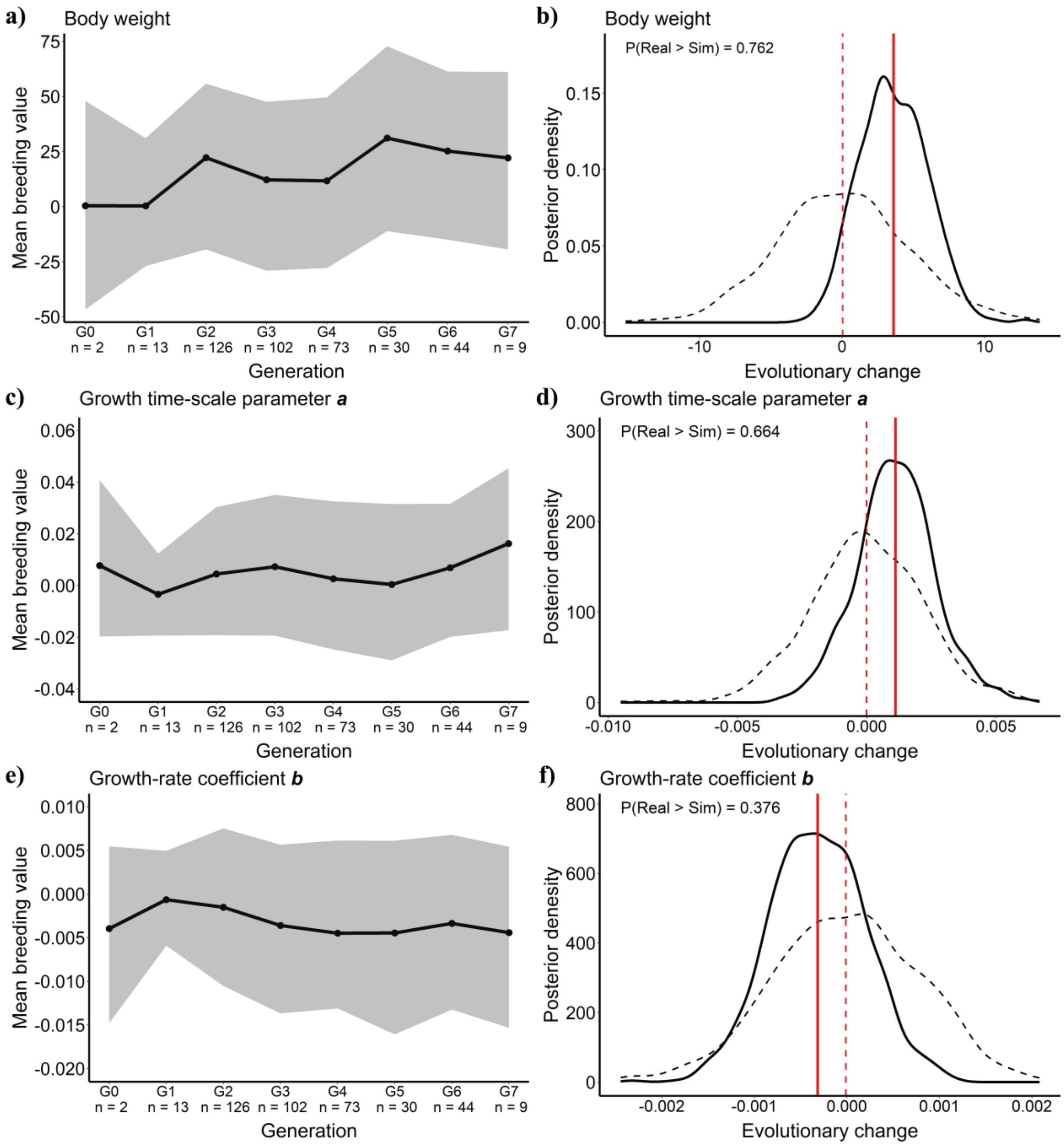
