## Supplementary figures and images for "Monitor individual health and improve breeding success in Crested ibis (*Nipponia nippon*)"

### Figure S3

Figure S3. The growth trend of all unhealthy individuals.

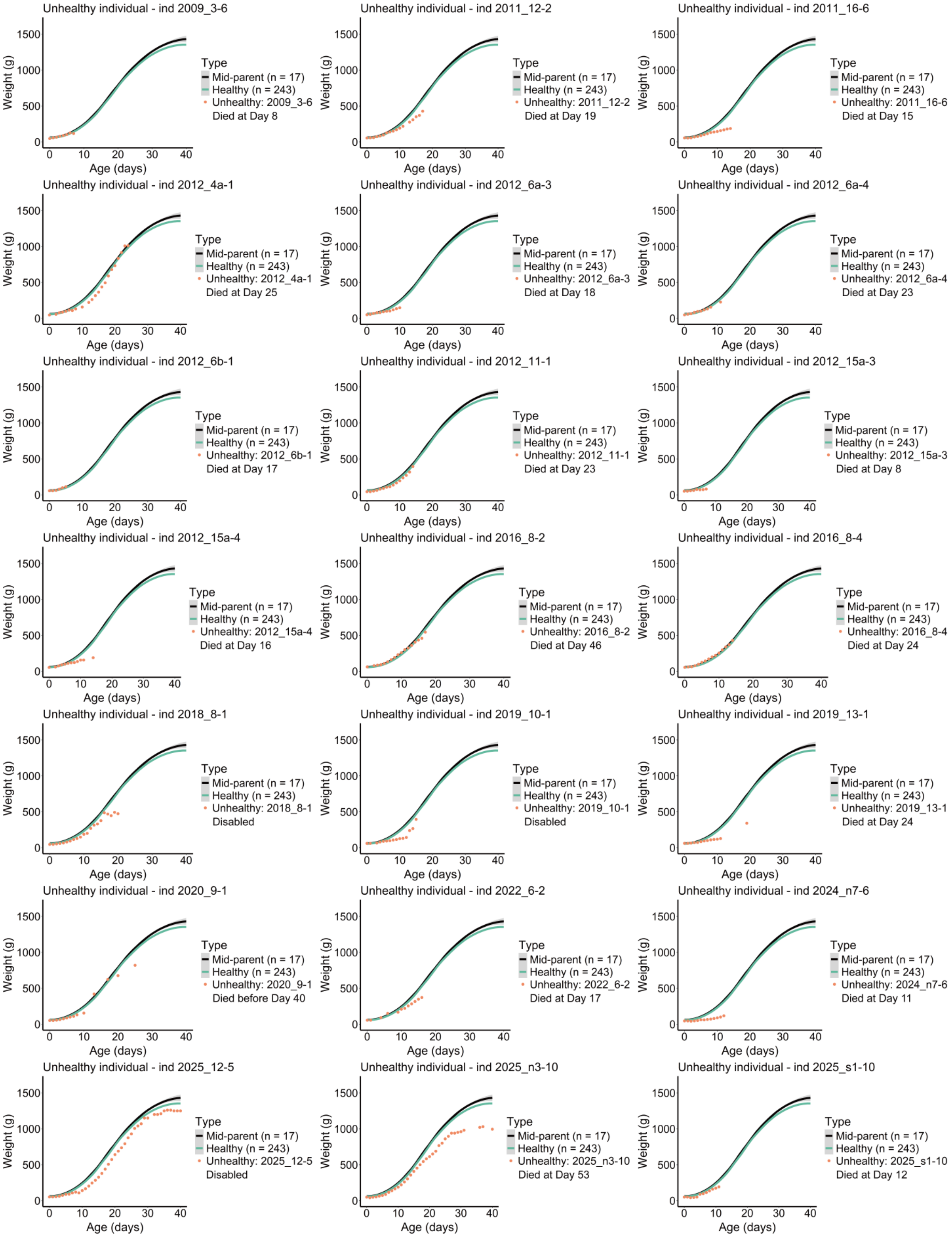
