## Supplementary material for "Monitor individual health and improve breeding success in Crested ibis (*Nipponia nippon*)": Table S1

Table S1. Estimated breeding values for body weight and growth curve parameters (growth time-scale parameter ***a*** and growth-rate coefficient ***b***) for all individuals.

| id | dam | sire | generation | Body weight (g) | growth time-scale parameter ***a*** | growth-rate coefficient ***b*** |
|---|---|---|---|---|---|---|
| Y | NA | NA | 0 | -0.165 | 0.00979 | -0.004022 |
| M | NA | NA | 0 | 0.979 | 0.00569 | -0.003920 |
| founder_1 | NA | NA | 1 | -22.118 | 0.00319 | -0.004349 |
| founder_3 | NA | NA | 1 | 44.019 | -0.01989 | -0.001288 |
| founder_5 | M | Y | 1 | -86.170 | 0.00521 | -0.011739 |
| founder_6 | M | Y | 1 | -62.221 | 0.03401 | -0.009076 |
| founder_7 | M | Y | 1 | -14.828 | 0.00140 | -0.003259 |
| founder_9 | M | Y | 1 | -7.895 | 0.00945 | -0.006466 |
| founder_10 | M | Y | 1 | -2.088 | 0.00917 | -0.004298 |
| founder_11 | M | Y | 1 | 58.231 | 0.00234 | -0.003058 |
| founder_12 | M | Y | 1 | 34.823 | 0.01597 | 0.000862 |
| founder_13 | M | Y | 1 | 20.212 | -0.00036 | -0.005154 |
| founder_14 | M | Y | 1 | -18.601 | 0.00835 | -0.004823 |
| founder_15 | M | Y | 1 | 32.444 | -0.00367 | -0.002469 |
| founder_17 | M | Y | 1 | 36.401 | 0.01429 | 0.000704 |
| 2008_2-1 | founder_13 | founder_3 | 2 | 106.950 | -0.01789 | -0.004498 |
| 2008_2-2 | founder_13 | founder_3 | 2 | -44.690 | -0.00986 | -0.008509 |
| 2008_2-3 | founder_13 | founder_3 | 2 | 43.062 | -0.01982 | -0.001351 |
| 2008_2-4 | founder_13 | founder_3 | 2 | -62.611 | -0.00080 | -0.008450 |
| 2008_2-5 | founder_13 | founder_3 | 2 | 23.176 | 0.01598 | -0.001858 |
| 2009_2-1 | founder_13 | founder_3 | 2 | -4.738 | -0.02581 | -0.007658 |
| 2009_2-2 | founder_13 | founder_3 | 2 | 1.201 | -0.00030 | -0.000895 |
| 2009_2-4 | founder_13 | founder_3 | 2 | 33.421 | -0.01372 | 0.001460 |
| 2009_2-5 | founder_13 | founder_3 | 2 | 40.916 | 0.00375 | -0.003574 |
| 2009_2-6 | founder_13 | founder_3 | 2 | 65.651 | -0.00834 | -0.003078 |
| 2009_4-3 | founder_9 | founder_1 | 2 | -3.772 | 0.01079 | -0.005981 |
| 2009_4-1 | founder_9 | founder_1 | 2 | -50.997 | 0.00661 | -0.009507 |
| 2009_3-3 | founder_15 | founder_10 | 2 | 22.198 | 0.00551 | -0.002240 |
| 2009_3-1 | founder_15 | founder_10 | 2 | 3.842 | -0.00846 | -0.003714 |
| 2009_3-2 | founder_15 | founder_10 | 2 | 41.121 | 0.00010 | -0.003815 |
| 2009_3-4 | founder_15 | founder_10 | 2 | 0.196 | 0.00955 | 0.001116 |
| 2009_3-5 | founder_15 | founder_10 | 2 | 14.112 | 0.01491 | -0.000419 |
| 2009_3-6 | founder_15 | founder_10 | 2 | 14.550 | 0.00205 | -0.003257 |
| 2009_1-1 | founder_17 | founder_12 | 2 | 28.550 | 0.00901 | 0.002308 |
| 2009_1-2 | founder_17 | founder_12 | 2 | 72.633 | 0.01880 | 0.001528 |
| 2009_1-4 | founder_17 | founder_12 | 2 | 60.750 | 0.00928 | -0.000122 |
| 2009_1-3 | founder_17 | founder_12 | 2 | 47.732 | 0.01501 | 0.002245 |
| 2009_1-5 | founder_17 | founder_12 | 2 | 84.810 | 0.02044 | 0.001922 |

| | | | | | | |
|---|---|---|---|---|---|---|
| 2010_7-4 | founder_13 | founder_3 | 2 | 46.500 | -0.00808 | -0.006363 |
| 2010_7-5 | founder_13 | founder_3 | 2 | 21.688 | -0.00877 | -0.005982 |
| 2010_3-1 | 2008_2-1 | founder_5 | 3 | 13.869 | -0.00816 | -0.009578 |
| 2010_3-3 | 2008_2-1 | founder_5 | 3 | 15.680 | -0.00987 | -0.006732 |
| 2010_3-2 | 2008_2-1 | founder_5 | 3 | 20.579 | -0.00119 | -0.005720 |
| 2010_3-4 | 2008_2-1 | founder_5 | 3 | -10.147 | -0.00159 | -0.011015 |
| 2010_3-5 | 2008_2-1 | founder_5 | 3 | -70.448 | -0.00730 | -0.011805 |
| 2010_5-1 | 2008_2-4 | founder_11 | 3 | 16.438 | -0.00822 | -0.004553 |
| 2010_5-4 | 2008_2-4 | founder_11 | 3 | -7.639 | 0.00078 | -0.007473 |
| 2010_5-5 | 2008_2-4 | founder_11 | 3 | 1.323 | 0.00610 | -0.009300 |
| 2010_6-2 | founder_15 | founder_10 | 2 | 46.228 | -0.01047 | -0.004130 |
| 2010_6-1 | founder_15 | founder_10 | 2 | 15.067 | 0.00215 | -0.006789 |
| 2010_6-4 | founder_15 | founder_10 | 2 | 21.394 | 0.00249 | -0.006666 |
| 2010_6-3 | founder_15 | founder_10 | 2 | 14.318 | 0.00202 | -0.005271 |
| 2010_6-5 | founder_15 | founder_10 | 2 | -18.223 | 0.00642 | -0.008258 |
| 2010_4-6 | 2008_2-5 | founder_6 | 3 | -47.153 | 0.02784 | -0.001607 |
| 2010_4-4 | 2008_2-5 | founder_6 | 3 | -22.971 | 0.03144 | -0.002079 |
| 2010_4-5 | 2008_2-5 | founder_6 | 3 | -44.283 | 0.03366 | -0.008086 |
| 2010_8-1 | founder_17 | founder_12 | 2 | 67.791 | 0.00001 | 0.007995 |
| 2010_8-2 | founder_17 | founder_12 | 2 | 63.817 | 0.01956 | 0.004832 |
| 2010_8-3 | founder_17 | founder_12 | 2 | -11.224 | 0.01644 | 0.004792 |
| 2011_7-1 | founder_13 | founder_3 | 2 | 44.591 | -0.01478 | -0.002037 |
| 2011_7-2 | founder_13 | founder_3 | 2 | 56.921 | -0.01527 | 0.001039 |
| 2011_7-3 | founder_13 | founder_3 | 2 | 40.181 | -0.02121 | -0.005440 |
| 2011_7-4 | founder_13 | founder_3 | 2 | 60.703 | -0.02920 | -0.006597 |
| 2011_7-5 | founder_13 | founder_3 | 2 | 52.937 | -0.00692 | 0.000104 |
| 2011_13-2 | 2009_3-3 | founder_14 | 3 | -27.995 | 0.00972 | -0.004566 |
| 2011_16-3 | 2009_1-3 | 2008_2-2 | 3 | 0.043 | 0.00082 | -0.004381 |
| 2011_16-1 | 2009_1-3 | 2008_2-2 | 3 | -3.892 | -0.00339 | -0.004865 |
| 2011_16-6 | 2009_1-3 | 2008_2-2 | 3 | -0.591 | 0.00281 | -0.003074 |
| 2011_17-1 | 2009_4-3 | 2009_3-2 | 3 | -13.328 | 0.01226 | -0.006349 |
| 2011_17-2 | 2009_4-3 | 2009_3-2 | 3 | -2.333 | 0.01271 | -0.005400 |
| 2011_15-1 | 2009_1-5 | 2009_4-1 | 3 | -27.458 | 0.01618 | -0.005175 |
| 2011_15-2 | 2009_1-5 | 2009_4-1 | 3 | 14.617 | 0.01675 | -0.003556 |
| 2011_15-5 | 2009_1-5 | 2009_4-1 | 3 | 26.592 | 0.00890 | -0.004657 |
| 2011_15-6 | 2009_1-5 | 2009_4-1 | 3 | 36.320 | 0.02086 | -0.002862 |
| 2011_15-7 | 2009_1-5 | 2009_4-1 | 3 | 24.118 | 0.00844 | -0.006217 |
| 2011_3-2 | 2008_2-1 | founder_5 | 3 | 21.399 | -0.00132 | -0.005802 |
| 2011_5-1 | 2008_2-4 | founder_11 | 3 | 16.499 | -0.00149 | -0.005066 |
| 2011_5-2 | 2008_2-4 | founder_11 | 3 | 6.675 | -0.00593 | -0.003978 |
| 2011_5-5 | 2008_2-4 | founder_11 | 3 | -2.850 | 0.00133 | -0.006035 |
| 2011_6-1 | founder_15 | founder_10 | 2 | -7.743 | 0.00960 | -0.005084 |

| | | | | | | |
|---|---|---|---|---|---|---|
| 2011_6-2 | founder_15 | founder_10 | 2 | 6.081 | 0.00462 | -0.003771 |
| 2011_6-3 | founder_15 | founder_10 | 2 | -0.512 | 0.00702 | -0.003581 |
| 2011_12-2 | 2009_2-2 | 2009_1-2 | 3 | 35.906 | 0.00977 | 0.000227 |
| 2011_12-5 | 2009_2-2 | 2009_1-2 | 3 | 42.776 | 0.01763 | 0.000757 |
| 2011_2-2 | 2009_3-1 | 2008_2-3 | 3 | 3.851 | -0.02015 | 0.001204 |
| 2011_2-1 | 2009_3-1 | 2008_2-3 | 3 | 42.293 | -0.01386 | -0.000213 |
| 2011_2-3 | 2009_3-1 | 2008_2-3 | 3 | 53.516 | -0.00808 | 0.002471 |
| 2011_2-4 | 2009_3-1 | 2008_2-3 | 3 | 47.969 | -0.02099 | -0.000546 |
| 2011_2-5 | 2009_3-1 | 2008_2-3 | 3 | 38.723 | -0.02053 | -0.005659 |
| 2011_4-4 | 2008_2-5 | founder_6 | 3 | -17.859 | 0.03529 | -0.006123 |
| 2011_4-1 | 2008_2-5 | founder_6 | 3 | -32.897 | 0.03041 | -0.005601 |
| 2011_4-2 | 2008_2-5 | founder_6 | 3 | -7.442 | 0.03226 | -0.004904 |
| 2011_4-5 | 2008_2-5 | founder_6 | 3 | -21.526 | 0.02557 | -0.005943 |
| 2011_14-1 | 2009_2-5 | founder_7 | 3 | -23.501 | -0.00158 | -0.004109 |
| 2011_14-2 | 2009_2-5 | founder_7 | 3 | -16.736 | 0.00068 | -0.005704 |
| 2011_14-3 | 2009_2-5 | founder_7 | 3 | -20.882 | 0.02501 | -0.002919 |
| 2011_8-1 | founder_17 | founder_12 | 2 | 34.910 | 0.01435 | 0.000648 |
| 2011_8-2 | founder_17 | founder_12 | 2 | 34.512 | 0.01540 | 0.000780 |
| 2011_8-3 | founder_17 | founder_12 | 2 | 32.458 | 0.01583 | 0.000773 |
| 2012_11-1 | 2010_6-1 | 2009_2-6 | 3 | 37.922 | -0.00289 | -0.005080 |
| 2012_12-1 | 2009_2-2 | 2009_1-2 | 3 | 36.460 | 0.00916 | 0.000271 |
| 2012_12-2 | 2009_2-2 | 2009_1-2 | 3 | 39.519 | 0.00938 | 0.000723 |
| 2012_12-3 | 2009_2-2 | 2009_1-2 | 3 | 37.411 | 0.00853 | 0.000317 |
| 2012_15a-2 | 2009_1-5 | 2009_4-1 | 3 | 68.590 | 0.01764 | -0.001977 |
| 2012_15a-1 | 2009_1-5 | 2009_4-1 | 3 | 59.326 | 0.01611 | 0.000823 |
| 2012_15a-4 | 2009_1-5 | 2009_4-1 | 3 | 18.427 | 0.01415 | -0.003782 |
| 2012_15a-3 | 2009_1-5 | 2009_4-1 | 3 | 16.627 | 0.01306 | -0.003442 |
| 2012_15b-2 | 2009_2-5 | founder_7 | 3 | 12.571 | 0.00299 | -0.003617 |
| 2012_15b-4 | 2009_2-5 | founder_7 | 3 | 15.182 | 0.00233 | -0.003551 |
| 2012_15b-5 | 2009_2-5 | founder_7 | 3 | 14.385 | 0.00381 | -0.003619 |
| 2012_17-1 | 2009_4-3 | 2009_3-2 | 3 | 63.297 | 0.00254 | -0.001516 |
| 2012_17-2 | 2009_4-3 | 2009_3-2 | 3 | 58.114 | -0.00130 | -0.002172 |
| 2012_18a-1 | 2010_3-3 | 2009_3-4 | 4 | 9.470 | -0.00029 | -0.002637 |
| 2012_18b-1 | NA | NA | NA | -17.560 | -0.02159 | -0.003686 |
| 2012_3a-1 | 2010_4-6 | 2010_3-5 | 4 | -60.488 | 0.00949 | -0.006617 |
| 2012_3a-3 | 2010_4-6 | 2010_3-5 | 4 | -59.768 | 0.00959 | -0.006776 |
| 2012_3a-2 | 2010_4-6 | 2010_3-5 | 4 | -55.692 | 0.01086 | -0.006564 |
| 2012_3b-2 | 2008_2-1 | founder_5 | 3 | 13.490 | -0.00652 | -0.008176 |
| 2012_3b-1 | 2008_2-1 | founder_5 | 3 | 7.078 | -0.00632 | -0.007832 |
| 2012_3b-3 | 2008_2-1 | founder_5 | 3 | -17.249 | -0.00828 | -0.010105 |
| 2012_3b-4 | 2008_2-1 | founder_5 | 3 | -46.787 | -0.01636 | -0.016926 |
| 2012_4a-1 | 2008_2-5 | founder_6 | 3 | -59.120 | 0.03016 | -0.006644 |

| | | | | | | |
|---|---|---|---|---|---|---|
| 2012_4a-5 | 2008_2-5 | founder_6 | 3 | -44.470 | 0.02773 | -0.008700 |
| 2012_4a-4 | 2008_2-5 | founder_6 | 3 | -75.280 | 0.03457 | -0.010440 |
| 2012_4b-1 | 2010_8-1 | 2010_4-5 | 4 | 28.961 | 0.01744 | 0.000684 |
| 2012_4b-2 | 2010_8-1 | 2010_4-5 | 4 | -17.365 | 0.01884 | -0.000482 |
| 2012_6a-1 | founder_15 | founder_10 | 2 | 57.568 | -0.00348 | 0.000290 |
| 2012_6a-2 | founder_15 | founder_10 | 2 | 19.474 | 0.00347 | -0.000876 |
| 2012_6a-3 | founder_15 | founder_10 | 2 | 14.595 | 0.00151 | -0.003002 |
| 2012_6a-4 | founder_15 | founder_10 | 2 | 17.275 | 0.00320 | -0.003778 |
| 2012_6b-1 | 2010_6-5 | 2010_5-5 | 4 | -6.861 | 0.00581 | -0.008719 |
| 2012_7-1 | founder_13 | founder_3 | 2 | 107.382 | -0.01626 | 0.001796 |
| 2012_7-2 | founder_13 | founder_3 | 2 | 22.150 | -0.00690 | -0.002262 |
| 2012_8-2 | founder_17 | founder_12 | 2 | 34.814 | 0.01495 | 0.000638 |
| 2012_8-1 | founder_17 | founder_12 | 2 | 38.593 | 0.01404 | 0.000426 |
| 2012_9-4 | 2010_8-2 | 2010_5-1 | 4 | -18.653 | 0.02200 | -0.003239 |
| 2013_15-2 | 2009_1-5 | 2009_3-4 | 3 | 43.331 | 0.01369 | 0.002008 |
| 2013_17-2 | 2009_4-3 | 2010_6-2 | 3 | 24.105 | 0.00048 | -0.005226 |
| 2013_17-3 | 2009_4-3 | 2010_6-2 | 3 | 21.143 | 0.00025 | -0.004953 |
| 2013_18-1 | 2010_3-3 | 2009_1-4 | 4 | 37.714 | -0.00173 | -0.003472 |
| 2013_18-2 | 2010_3-3 | 2009_1-4 | 4 | 38.389 | -0.00075 | -0.003005 |
| 2013_3-2 | 2008_2-1 | founder_5 | 3 | 12.446 | -0.00676 | -0.008538 |
| 2013_5-2 | 2008_2-4 | founder_11 | 3 | -0.597 | 0.00063 | -0.005394 |
| 2013_5-3 | 2008_2-4 | founder_11 | 3 | -1.731 | 0.00062 | -0.005555 |
| 2013_5-1 | 2008_2-4 | founder_11 | 3 | -0.939 | 0.00173 | -0.005468 |
| 2013_6-1 | founder_15 | founder_7 | 2 | 8.396 | -0.00293 | -0.002911 |
| 2013_6-2 | founder_15 | founder_7 | 2 | 10.162 | -0.00044 | -0.003073 |
| 2013_6-4 | founder_15 | founder_7 | 2 | 6.901 | -0.00085 | -0.003094 |
| 2013_6-3 | founder_15 | founder_7 | 2 | 8.812 | -0.00107 | -0.002918 |
| 2013_7-1 | founder_13 | founder_3 | 2 | 33.273 | -0.01074 | -0.003139 |
| 2013_7-3 | founder_13 | founder_3 | 2 | 33.081 | -0.00921 | -0.003708 |
| 2013_8-1 | founder_17 | founder_12 | 2 | 34.109 | 0.01425 | 0.000720 |
| 2013_8-4 | founder_17 | founder_12 | 2 | 14.531 | 0.02633 | 0.001218 |
| 2013_8-2 | founder_17 | founder_12 | 2 | 39.160 | 0.01531 | 0.000686 |
| 2013_8-3 | founder_17 | founder_12 | 2 | 37.982 | 0.01477 | 0.000770 |
| 2013_8-5 | founder_17 | founder_12 | 2 | 33.782 | 0.01790 | -0.003120 |
| 2013_8-6 | founder_17 | founder_12 | 2 | 35.725 | 0.01614 | 0.001303 |
| 2013_9-4 | 2010_8-2 | 2010_5-1 | 4 | 39.120 | 0.00764 | -0.002991 |
| 2013_9-3 | 2010_8-2 | 2010_5-1 | 4 | 39.996 | 0.00513 | -0.000051 |
| 2013_9-2 | 2010_8-2 | 2010_5-1 | 4 | 42.517 | 0.00234 | 0.002822 |
| 2013_9-1 | 2010_8-2 | 2010_5-1 | 4 | 42.118 | 0.00644 | -0.000065 |
| 2013_9-5 | 2010_8-2 | 2010_5-1 | 4 | 38.984 | 0.00538 | 0.000128 |
| 2014_7-1 | founder_13 | founder_3 | 2 | 34.906 | -0.01525 | -0.000059 |
| 2014_7-2 | founder_13 | founder_3 | 2 | 32.033 | -0.00963 | -0.003116 |

| | | | | | | |
|---|---|---|---|---|---|---|
| 2014_2-3 | founder_9 | 2010_5-4 | 4 | -6.859 | 0.00564 | -0.007076 |
| 2014_12-1 | 2010_6-5 | 2010_5-5 | 4 | -9.270 | 0.00640 | -0.008633 |
| 2014_12-3 | 2010_6-5 | 2010_5-5 | 4 | -7.816 | 0.00577 | -0.008588 |
| 2014_12-2 | 2010_6-5 | 2010_5-5 | 4 | -8.647 | 0.00656 | -0.009063 |
| 2014_12-5 | 2010_6-5 | 2010_5-5 | 4 | -8.968 | 0.00627 | -0.009055 |
| 2014_9-1 | 2010_8-2 | 2010_5-1 | 4 | 40.589 | 0.00698 | 0.000091 |
| 2014_1-3 | 2011_15-7 | 2011_7-3 | 4 | 40.327 | 0.00510 | -0.005389 |
| 2014_1-2 | 2011_15-7 | 2011_7-3 | 4 | 9.538 | 0.00396 | -0.004064 |
| 2014_1-4 | 2011_15-7 | 2011_7-3 | 4 | 32.403 | -0.00658 | -0.005898 |
| 2014_3-1 | 2008_2-1 | founder_5 | 3 | 22.952 | -0.00725 | -0.007387 |
| 2014_3-2 | 2008_2-1 | founder_5 | 3 | 26.138 | -0.00494 | -0.007995 |
| 2014_3-3 | 2008_2-1 | founder_5 | 3 | 7.536 | -0.00533 | -0.007372 |
| 2014_3-4 | 2008_2-1 | founder_5 | 3 | 11.903 | -0.00702 | -0.008214 |
| 2014_3-5 | 2008_2-1 | founder_5 | 3 | 10.189 | -0.00649 | -0.007965 |
| 2014_5-1 | 2008_2-4 | founder_11 | 3 | -4.933 | 0.00047 | -0.005792 |
| 2014_5-2 | 2008_2-4 | founder_11 | 3 | 6.239 | 0.01029 | -0.000358 |
| 2014_5-3 | 2008_2-4 | founder_11 | 3 | -1.057 | 0.00209 | -0.005549 |
| 2014_5-4 | 2008_2-4 | founder_11 | 3 | -4.838 | 0.00263 | -0.005977 |
| 2014_13-1 | 2010_3-3 | 2009_1-4 | 4 | 28.411 | -0.01231 | -0.005698 |
| 2014_13-2 | 2010_3-3 | 2009_1-4 | 4 | 34.258 | -0.00145 | -0.004238 |
| 2014_13-4 | 2010_3-3 | 2009_1-4 | 4 | 36.766 | 0.00047 | -0.003546 |
| 2014_13-3 | 2010_3-3 | 2009_1-4 | 4 | 37.780 | -0.00032 | -0.003626 |
| 2014_13-7 | 2010_3-3 | 2009_1-4 | 4 | 41.194 | -0.00008 | -0.003168 |
| 2014_13-5 | 2010_3-3 | 2009_1-4 | 4 | 39.681 | 0.00029 | -0.003367 |
| 2014_13-6 | 2010_3-3 | 2009_1-4 | 4 | 39.599 | 0.00076 | -0.003047 |
| 2014_4-1 | 2008_2-5 | founder_6 | 3 | -8.358 | 0.02515 | -0.005736 |
| 2014_4-3 | 2008_2-5 | founder_6 | 3 | -3.796 | 0.02604 | -0.003871 |
| 2014_4-2 | 2008_2-5 | founder_6 | 3 | -22.186 | 0.02373 | -0.005052 |
| 2014_4-4 | 2008_2-5 | founder_6 | 3 | -3.308 | 0.02291 | -0.007422 |
| 2014_8-1 | founder_17 | founder_12 | 2 | 36.398 | 0.01563 | 0.000949 |
| 2014_8-2 | founder_17 | founder_12 | 2 | 36.877 | 0.01602 | 0.000392 |
| 2014_8-3 | founder_17 | founder_12 | 2 | 28.782 | 0.01913 | -0.001272 |
| 2014_8-4 | founder_17 | founder_12 | 2 | 36.173 | 0.01454 | 0.000915 |
| 2015_2-3 | founder_9 | 2010_5-4 | 4 | -6.630 | 0.00612 | -0.007041 |
| 2015_12-3 | 2010_6-5 | 2010_5-5 | 4 | -8.170 | 0.00423 | -0.009189 |
| 2015_12-1 | 2010_6-5 | 2010_5-5 | 4 | -0.958 | 0.00688 | -0.015599 |
| 2015_12-2 | 2010_6-5 | 2010_5-5 | 4 | -10.413 | 0.00664 | -0.009016 |
| 2015_9-2 | 2010_8-2 | 2010_5-1 | 4 | 39.974 | 0.00578 | 0.000169 |
| 2015_9-3 | 2010_8-2 | 2010_5-1 | 4 | 40.378 | 0.00622 | 0.000211 |
| 2015_16-1 | 2013_9-4 | 2013_8-5 | 5 | 48.237 | 0.01435 | -0.003091 |
| 2015_16-4 | 2013_9-4 | 2013_8-5 | 5 | 34.189 | 0.00836 | -0.003343 |
| 2015_16-3 | 2013_9-4 | 2013_8-5 | 5 | 36.677 | 0.01279 | -0.003244 |

| | | | | | | |
|---|---|---|---|---|---|---|
| 2015_3-3 | 2008_2-1 | founder_5 | 3 | 10.115 | -0.00608 | -0.008050 |
| 2015_3-4 | 2008_2-1 | founder_5 | 3 | 12.355 | -0.00605 | -0.008376 |
| 2015_5-2 | 2008_2-4 | founder_11 | 3 | -0.926 | 0.00153 | -0.005609 |
| 2015_5-1 | 2008_2-4 | founder_11 | 3 | -1.142 | 0.00081 | -0.006064 |
| 2015_5-3 | 2008_2-4 | founder_11 | 3 | 40.141 | -0.00013 | -0.002771 |
| 2015_13-2 | 2010_3-3 | 2009_1-4 | 4 | 39.402 | -0.00041 | -0.003369 |
| 2015_13-5 | 2010_3-3 | 2009_1-4 | 4 | 38.470 | -0.00126 | -0.003637 |
| 2015_14-2 | 2008_2-5 | founder_6 | 3 | -10.879 | 0.01750 | -0.011600 |
| 2015_6-1 | founder_15 | founder_7 | 2 | -21.927 | -0.00401 | -0.006002 |
| 2015_6-2 | founder_15 | founder_7 | 2 | -3.658 | -0.00703 | 0.000886 |
| 2015_6-3 | founder_15 | founder_7 | 2 | 1.791 | -0.00984 | -0.004730 |
| 2015_8-1 | founder_17 | founder_12 | 2 | 34.551 | 0.01431 | 0.000428 |
| 2015_8-2 | founder_17 | founder_12 | 2 | 35.418 | 0.01334 | 0.000577 |
| 2015_8-7 | founder_17 | founder_12 | 2 | 35.673 | 0.01485 | 0.000651 |
| 2015_8-6 | founder_17 | founder_12 | 2 | 36.056 | 0.01536 | 0.000634 |
| 2015_8-5 | founder_17 | founder_12 | 2 | 36.013 | 0.01550 | 0.000954 |
| 2015_8-4 | founder_17 | founder_12 | 2 | 36.709 | 0.01493 | -0.000025 |
| 2016_2-4 | founder_9 | 2010_5-4 | 4 | -7.358 | 0.00570 | -0.006824 |
| 2016_9-2 | 2010_8-2 | 2010_5-1 | 4 | 45.127 | 0.00629 | 0.002263 |
| 2016_1-1 | 2012_15a-1 | 2011_7-3 | 4 | 23.057 | 0.00088 | 0.002272 |
| 2016_16-2 | 2013_9-4 | 2013_8-5 | 5 | 36.167 | 0.01332 | -0.003386 |
| 2016_16-3 | 2013_9-4 | 2013_8-5 | 5 | 36.500 | 0.01227 | -0.002766 |
| 2016_16-5 | 2013_9-4 | 2013_8-5 | 5 | 22.795 | 0.01359 | 0.001071 |
| 2016_16-6 | 2013_9-4 | 2013_8-5 | 5 | 31.295 | 0.00975 | -0.006345 |
| 2016_16-8 | 2013_9-4 | 2013_8-5 | 5 | 34.910 | 0.01185 | -0.003355 |
| 2016_16-9 | 2013_9-4 | 2013_8-5 | 5 | 37.815 | 0.01276 | -0.002903 |
| 2016_4-1 | 2014_3-2 | 2014_3-1 | 4 | 25.713 | -0.00103 | -0.010161 |
| 2016_5-2 | 2008_2-4 | founder_11 | 3 | 18.120 | -0.00609 | -0.009347 |
| 2016_13-1 | 2010_3-3 | 2009_1-4 | 4 | 38.088 | -0.00067 | -0.002991 |
| 2016_13-2 | 2010_3-3 | 2009_1-4 | 4 | 17.364 | 0.00740 | -0.004833 |
| 2016_13-7 | 2010_3-3 | 2009_1-4 | 4 | 37.345 | 0.00340 | -0.002445 |
| 2016_13-6 | 2010_3-3 | 2009_1-4 | 4 | 29.847 | -0.01670 | -0.003915 |
| 2016_13-5 | 2010_3-3 | 2009_1-4 | 4 | 39.456 | 0.00055 | -0.003205 |
| 2016_14-4 | 2008_2-5 | founder_6 | 3 | -22.542 | 0.02493 | -0.005585 |
| 2016_6-3 | founder_15 | founder_7 | 2 | 8.657 | -0.00175 | -0.002642 |
| 2016_6-4 | founder_15 | founder_7 | 2 | 45.658 | 0.00288 | -0.006072 |
| 2016_6-8 | founder_15 | founder_7 | 2 | 6.449 | -0.00085 | -0.002571 |
| 2016_8-1 | founder_17 | founder_12 | 2 | 37.009 | 0.01575 | 0.000456 |
| 2016_8-3 | founder_17 | founder_12 | 2 | 66.083 | 0.01695 | -0.003504 |
| 2016_8-4 | founder_17 | founder_12 | 2 | 36.897 | 0.01334 | 0.000904 |
| 2016_8-2 | founder_17 | founder_12 | 2 | 34.067 | 0.01505 | 0.000637 |
| 2016_8-5 | founder_17 | founder_12 | 2 | 33.237 | 0.01508 | 0.000775 |

| | | | | | | |
|---|---|---|---|---|---|---|
| 2016_8-6 | founder_17 | founder_12 | 2 | 35.347 | 0.01539 | 0.000388 |
| 2016_8-7 | founder_17 | founder_12 | 2 | 35.320 | 0.01597 | 0.000840 |
| 2016_NA-1 | NA | NA | NA | 1.201 | -0.00086 | -0.000152 |
| 2017_15-1 | 2012_15a-1 | 2011_7-3 | 4 | 47.500 | -0.00375 | -0.002533 |
| 2017_5-1 | 2015_16-1 | 2014_4-1 | 6 | 22.127 | 0.02017 | -0.004368 |
| 2017_5-2 | 2015_16-1 | 2014_4-1 | 6 | 19.460 | 0.01851 | -0.004408 |
| 2017_4-1 | 2014_3-2 | 2014_3-1 | 4 | 24.287 | -0.00734 | -0.008011 |
| 2017_4-2 | 2014_3-2 | 2014_3-1 | 4 | 23.284 | -0.00621 | -0.007423 |
| 2017_4-3 | 2014_3-2 | 2014_3-1 | 4 | 22.035 | -0.00530 | -0.007693 |
| 2017_luo-4 | 2008_2-1 | founder_5 | 3 | -27.820 | -0.00985 | -0.011521 |
| 2017_luo-1 | 2008_2-1 | founder_5 | 3 | 12.018 | -0.00724 | -0.007934 |
| 2017_luo-3 | 2008_2-1 | founder_5 | 3 | 9.768 | -0.00553 | -0.007532 |
| 2017_luo-5 | 2008_2-1 | founder_5 | 3 | 9.235 | -0.00670 | -0.008302 |
| 2017_luo-6 | 2008_2-1 | founder_5 | 3 | 10.687 | -0.00566 | -0.008562 |
| 2017_13-1 | 2010_3-3 | 2009_1-4 | 4 | 36.869 | -0.00040 | -0.003337 |
| 2017_13-2 | 2010_3-3 | 2009_1-4 | 4 | 69.446 | -0.00018 | -0.000960 |
| 2017_13-3 | 2010_3-3 | 2009_1-4 | 4 | 37.111 | 0.00057 | -0.003429 |
| 2017_13-4 | 2010_3-3 | 2009_1-4 | 4 | 37.740 | -0.00040 | -0.003566 |
| 2017_13-5 | 2010_3-3 | 2009_1-4 | 4 | -5.524 | -0.00627 | -0.009857 |
| 2017_6-2 | founder_15 | founder_7 | 2 | 10.434 | -0.00107 | -0.002911 |
| 2017_6-3 | founder_15 | founder_7 | 2 | 12.348 | -0.00212 | -0.002686 |
| 2017_6-1 | founder_15 | founder_7 | 2 | 9.973 | -0.00069 | -0.002362 |
| 2017_6-4 | founder_15 | founder_7 | 2 | 7.194 | -0.00115 | -0.002946 |
| 2017_8-2 | founder_17 | founder_12 | 2 | 36.563 | 0.01531 | 0.001398 |
| 2017_8-3 | founder_17 | founder_12 | 2 | 34.848 | 0.01486 | 0.000882 |
| 2017_8-4 | founder_17 | founder_12 | 2 | 33.399 | 0.01369 | 0.000317 |
| 2017_8-1 | founder_17 | founder_12 | 2 | 37.329 | 0.01585 | 0.000745 |
| 2017_8-6 | founder_17 | founder_12 | 2 | 35.107 | 0.01527 | 0.000524 |
| 2017_8-5 | founder_17 | founder_12 | 2 | 34.956 | 0.01522 | 0.000763 |
| 2018_3-1 | 2014_1-3 | 2012_4a-5 | 5 | 19.081 | 0.02033 | -0.008287 |
| 2018_16-1 | 2014_1-2 | 2013_8-4 | 5 | 1.878 | 0.01558 | 0.000867 |
| 2018_14-2 | 2015_16-1 | 2014_4-1 | 6 | 33.342 | 0.01605 | -0.002644 |
| 2018_5-2 | 2011_15-7 | 2014_3-1 | 4 | 23.633 | -0.00015 | -0.006327 |
| 2018_4-1 | 2014_3-2 | 2014_3-1 | 4 | 24.394 | -0.00518 | -0.007674 |
| 2018_8-1 | 2015_6-1 | 2015_6-2 | 3 | -12.018 | -0.00555 | -0.002491 |
| 2018_8-3 | 2015_6-1 | 2015_6-2 | 3 | -19.276 | -0.00641 | -0.001484 |
| 2018_8-2 | 2015_6-1 | 2015_6-2 | 3 | -12.558 | -0.00568 | -0.002724 |
| 2018_6-1 | 2010_3-3 | 2009_1-4 | 4 | 23.576 | -0.00402 | -0.001588 |
| 2018_15-6 | founder_15 | founder_7 | 2 | 92.945 | -0.00981 | 0.003375 |
| 2018_9-1 | founder_17 | founder_12 | 2 | -5.782 | 0.00550 | 0.000173 |
| 2018_9-3 | founder_17 | founder_12 | 2 | -25.453 | 0.02762 | -0.006638 |
| 2018_9-4 | founder_17 | founder_12 | 2 | 33.708 | 0.01519 | 0.001229 |

| | | | | | | |
|---|---|---|---|---|---|---|
| 2018_9-5 | founder_17 | founder_12 | 2 | 14.683 | 0.01154 | -0.002372 |
| 2018_9-6 | founder_17 | founder_12 | 2 | 51.556 | 0.01301 | -0.000009 |
| 2018_luo-2 | NA | NA | NA | -1.580 | 0.00041 | 0.000007 |
| 2018_luo-4 | NA | NA | NA | 1.294 | -0.00021 | 0.000156 |
| 2018_luo-5 | NA | NA | NA | -1.702 | -0.00088 | 0.000036 |
| 2018_luo-6 | NA | NA | NA | 2.700 | -0.00188 | -0.000312 |
| 2019_4-1 | 2015_12-1 | 2015_16-4 | 6 | 11.605 | 0.00104 | -0.010518 |
| 2019_6-1 | 2010_3-3 | 2009_1-4 | 4 | 9.053 | -0.00009 | -0.003434 |
| 2019_6-2 | 2010_3-3 | 2009_1-4 | 4 | -17.976 | -0.00044 | -0.003102 |
| 2019_6-3 | 2010_3-3 | 2009_1-4 | 4 | 19.352 | 0.00034 | -0.003536 |
| 2019_9-1 | founder_17 | founder_12 | 2 | 43.838 | 0.00255 | -0.000993 |
| 2019_9-3 | founder_17 | founder_12 | 2 | 38.134 | 0.01491 | 0.000982 |
| 2019_9-2 | founder_17 | founder_12 | 2 | 34.887 | 0.01574 | 0.000460 |
| 2019_10-1 | NA | NA | NA | -8.077 | -0.00195 | 0.000531 |
| 2019_10-2 | NA | NA | NA | 6.048 | -0.00014 | 0.000203 |
| 2019_10-4 | NA | NA | NA | -2.168 | 0.00134 | -0.000101 |
| 2019_13-1 | 2015_13-2 | 2016_16-5 | 6 | 27.621 | 0.00654 | -0.001372 |
| 2019_14-1 | 2015_16-1 | 2014_4-1 | 6 | 33.815 | 0.02129 | -0.008378 |
| 2019_14-2 | 2015_16-1 | 2014_4-1 | 6 | 35.594 | 0.01624 | -0.005374 |
| 2019_14-3 | 2015_16-1 | 2014_4-1 | 6 | 28.871 | 0.01436 | -0.008184 |
| 2019_14-4 | 2015_16-1 | 2014_4-1 | 6 | 22.259 | 0.01987 | -0.004341 |
| 2019_15-1 | founder_15 | founder_7 | 2 | 11.052 | -0.00019 | -0.002743 |
| 2019_17-3 | 2013_9-4 | 2013_8-5 | 5 | 39.108 | 0.01201 | -0.003145 |
| 2019_luo1-1 | NA | NA | NA | -0.015 | -0.00120 | -0.000175 |
| 2019_luo1-2 | NA | NA | NA | 2.310 | -0.00043 | 0.000663 |
| 2019_luo1-4 | NA | NA | NA | -11.873 | -0.00112 | -0.000234 |
| 2019_luo2-1 | NA | NA | NA | -29.570 | 0.00114 | 0.000323 |
| 2019_luo2-3 | NA | NA | NA | 18.467 | -0.00135 | -0.000793 |
| 2020_13-1 | 2014_7-1 | 2013_9-2 | 5 | 41.412 | -0.01562 | 0.007340 |
| 2020_16-1 | 2013_9-4 | 2013_8-5 | 5 | 45.323 | 0.02142 | -0.008647 |
| 2020_5-2 | 2011_15-7 | 2014_3-1 | 4 | 13.530 | -0.00322 | -0.005753 |
| 2020_11-1 | 2015_12-1 | 2015_16-4 | 6 | 16.276 | 0.00734 | -0.009349 |
| 2020_6-8 | 2010_3-3 | 2009_1-4 | 4 | 36.788 | 0.00016 | -0.003946 |
| 2020_9-1 | founder_17 | founder_12 | 2 | 35.693 | 0.01478 | 0.000889 |
| 2020_9-2 | founder_17 | founder_12 | 2 | 43.767 | 0.02604 | 0.006221 |
| 2020_2-1 | NA | NA | NA | -2.186 | -0.00112 | -0.000423 |
| 2020_first-1 | NA | NA | NA | 26.099 | 0.00372 | 0.000188 |
| 2021_14-1 | 2015_16-1 | 2014_4-1 | 6 | 20.521 | 0.01891 | -0.004747 |
| 2021_5z-1 | 2011_15-7 | 2014_3-1 | 4 | 23.616 | -0.00023 | -0.007070 |
| 2021_5z-4 | 2011_15-7 | 2014_3-1 | 4 | 22.449 | -0.00002 | -0.006743 |
| 2021_4z-2 | 2014_3-2 | 2014_3-1 | 4 | 18.007 | -0.00863 | -0.008422 |
| 2021_4z-3 | 2014_3-2 | 2014_3-1 | 4 | 61.315 | -0.00659 | -0.004439 |

| | | | | | | |
|---|---|---|---|---|---|---|
| 2021_4z-1 | 2014_3-2 | 2014_3-1 | 4 | 26.106 | -0.00623 | -0.007581 |
| 2021_9-2 | founder_17 | founder_12 | 2 | -14.228 | 0.01263 | -0.003231 |
| 2021_9-1 | founder_17 | founder_12 | 2 | 42.524 | 0.01994 | 0.000898 |
| 2021_11z-2 | NA | NA | NA | 1.396 | 0.00059 | 0.000077 |
| 2021_11z-3 | NA | NA | NA | 2.988 | 0.00018 | 0.000044 |
| 2021_11z-1 | NA | NA | NA | -3.240 | 0.00027 | 0.000533 |
| 2021_11y-4 | NA | NA | NA | -3.525 | 0.00017 | -0.000179 |
| unknown1 | NA | NA | NA | 1.165 | 0.00071 | -0.000295 |
| unknown2 | NA | NA | NA | -20.086 | -0.00771 | 0.002756 |
| 2022_17-1 | 2014_1-2 | 2013_8-4 | 5 | 8.004 | 0.03320 | -0.001410 |
| 2022_17-2b | 2014_1-2 | 2013_8-4 | 5 | -8.564 | 0.01452 | -0.002424 |
| 2022_4-1 | 2015_5-3 | 2017_13-2 | 5 | 128.559 | 0.00985 | 0.000441 |
| 2022_16-5 | 2013_9-4 | 2013_8-5 | 5 | 35.297 | 0.01206 | -0.003231 |
| 2022_14-1 | 2015_16-1 | 2014_4-1 | 6 | -61.177 | 0.02428 | 0.001552 |
| 2022_14-4 | 2015_16-1 | 2014_4-1 | 6 | 38.787 | 0.02494 | -0.006251 |
| 2022_10-4 | 2017_13-5 | 2019_14-3 | 7 | -41.881 | 0.00195 | -0.015605 |
| 2022_10-6 | 2017_13-5 | 2019_14-3 | 7 | -19.510 | -0.00448 | -0.014686 |
| 2022_3-4 | 2015_12-1 | 2018_9-3 | 5 | 5.098 | 0.01246 | -0.012834 |
| 2022_3-5 | 2015_12-1 | 2018_9-3 | 5 | -47.750 | 0.02441 | -0.011267 |
| 2022_2-3 | 2020_first-1 | 2020_16-1 | NA | 4.524 | 0.02545 | -0.003473 |
| 2022_2-4 | 2020_first-1 | 2020_16-1 | NA | 89.277 | 0.00168 | -0.005088 |
| 2022_6-1 | 2010_3-3 | 2009_1-4 | 4 | 113.598 | -0.00271 | 0.001627 |
| 2022_6-2 | 2010_3-3 | 2009_1-4 | 4 | 38.637 | -0.00007 | -0.003623 |
| 2022_9-2 | founder_17 | founder_12 | 2 | 35.425 | 0.01484 | 0.001141 |
| 2022_9-3 | founder_17 | founder_12 | 2 | 38.242 | 0.01399 | 0.001279 |
| 2022_wild_AV | unknown2 | 2020_5-2 | NA | -2.609 | -0.00501 | -0.001715 |
| 2022_1-3 | NA | NA | NA | -29.156 | 0.01474 | 0.006922 |
| 2022_18-2 | NA | NA | NA | 39.841 | 0.00506 | 0.008499 |
| 2023_n7-2 | 2016_16-6 | 2010_5-4 | 6 | 11.836 | 0.00626 | -0.007091 |
| 2023_n7-3 | 2016_16-6 | 2010_5-4 | 6 | 12.288 | 0.00563 | -0.006532 |
| 2023_14-1 | 2015_16-1 | 2014_4-1 | 6 | 20.225 | 0.02137 | -0.004613 |
| 2023_14-2 | 2015_16-1 | 2014_4-1 | 6 | 63.701 | 0.01559 | -0.003808 |
| 2023_14-4 | 2015_16-1 | 2014_4-1 | 6 | 19.587 | 0.02057 | -0.004564 |
| 2023_14-5 | 2015_16-1 | 2014_4-1 | 6 | 21.698 | 0.01981 | -0.004483 |
| 2023_3-1 | 2015_12-1 | 2016_16-5 | 6 | 9.796 | 0.01012 | -0.007943 |
| 2023_3-2 | 2015_12-1 | 2016_16-5 | 6 | 12.533 | 0.01096 | -0.007043 |
| 2023_3-3 | 2015_12-1 | 2016_16-5 | 6 | 12.737 | 0.01082 | -0.007571 |
| 2023_3-4 | 2015_12-1 | 2016_16-5 | 6 | 11.380 | 0.01120 | -0.007383 |
| 2023_n5-2 | 2019_14-1 | 2019_6-2 | 7 | 8.537 | 0.01121 | -0.006385 |
| 2023_n5-3 | 2019_14-1 | 2019_6-2 | 7 | 7.552 | 0.01027 | -0.005694 |
| 2023_16-1 | 2019_14-2 | 2018_14-2 | 7 | 28.195 | 0.00875 | -0.002214 |
| 2023_16-3 | 2019_14-2 | 2018_14-2 | 7 | -16.766 | 0.00740 | -0.004226 |

| | | | | | | |
|---|---|---|---|---|---|---|
| 2023_16-4 | 2019_14-2 | 2018_14-2 | 7 | 82.961 | 0.01295 | -0.001414 |
| 2023_n6-1 | unknown2 | 2020_5-2 | NA | -3.720 | -0.00541 | -0.001849 |
| 2023_n6-2 | unknown2 | 2020_5-2 | NA | -2.055 | -0.00501 | -0.001905 |
| 2023_n8-2 | 2010_3-3 | 2009_1-4 | 4 | 37.526 | -0.00028 | -0.003283 |
| 2023_15-4 | founder_15 | founder_7 | 2 | -57.379 | -0.00247 | -0.003467 |
| 2023_15-6 | founder_15 | founder_7 | 2 | 11.746 | -0.00228 | -0.003196 |
| 2023_9-1 | founder_17 | founder_12 | 2 | -37.287 | 0.00198 | -0.004295 |
| 2023_9-2 | founder_17 | founder_12 | 2 | 33.151 | 0.01413 | 0.000718 |
| 2023_9-4 | founder_17 | founder_12 | 2 | 98.813 | 0.06669 | 0.003872 |
| 2023_9-6 | founder_17 | founder_12 | 2 | 94.494 | 0.00468 | 0.006619 |
| 2024_3-4 | 2015_12-1 | 2016_16-5 | 6 | 8.496 | 0.01057 | -0.007631 |
| 2024_3-5 | 2015_12-1 | 2016_16-5 | 6 | 8.985 | 0.01032 | -0.007090 |
| 2024_3-7 | 2015_12-1 | 2016_16-5 | 6 | 13.204 | 0.00937 | -0.006997 |
| 2024_6-2 | 2015_5-3 | 2017_13-2 | 5 | 54.880 | -0.00051 | -0.002192 |
| 2024_6-7 | 2015_5-3 | 2017_13-2 | 5 | 116.406 | 0.01595 | 0.000503 |
| 2024_9-1 | founder_17 | founder_12 | 2 | -26.126 | 0.00549 | 0.002241 |
| 2024_9-2 | founder_17 | founder_12 | 2 | 33.224 | 0.01499 | 0.001161 |
| 2024_9-3 | founder_17 | founder_12 | 2 | 33.824 | 0.01438 | 0.000692 |
| 2024_14-2 | 2015_16-1 | 2014_4-1 | 6 | 19.367 | 0.01924 | -0.004468 |
| 2024_14-3 | 2015_16-1 | 2014_4-1 | 6 | 7.377 | 0.00566 | -0.009500 |
| 2024_14-5 | 2015_16-1 | 2014_4-1 | 6 | 44.865 | 0.02052 | 0.001008 |
| 2024_14-6 | 2015_16-1 | 2014_4-1 | 6 | 23.111 | 0.01868 | -0.004566 |
| 2024_14-8 | 2015_16-1 | 2014_4-1 | 6 | 20.467 | 0.02055 | -0.004202 |
| 2024_14-9 | 2015_16-1 | 2014_4-1 | 6 | 37.821 | 0.02016 | -0.002751 |
| 2024_16-5 | 2014_1-2 | 2013_8-4 | 5 | 12.707 | 0.01618 | -0.001524 |
| 2024_18-1 | 2016_16-6 | 2010_5-4 | 6 | 14.154 | 0.00443 | -0.006923 |
| 2024_n3-1 | 2019_4-1 | 2019_9-1 | 7 | 24.308 | 0.00207 | -0.009215 |
| 2024_n5-7 | 2022_2-3 | 2022_4-1 | NA | 64.253 | 0.01779 | -0.001703 |
| 2024_n6-1 | unknown2 | 2020_5-2 | NA | -2.817 | -0.00607 | -0.001460 |
| 2024_n6-2 | unknown2 | 2020_5-2 | NA | -1.698 | -0.00566 | -0.001404 |
| 2024_n6-3 | unknown2 | 2020_5-2 | NA | -1.366 | -0.00605 | -0.001732 |
| 2024_n7-1 | 2022_3-5 | 2022_17-1 | 6 | -20.767 | 0.02857 | -0.006561 |
| 2024_n7-3 | 2022_3-5 | 2022_17-1 | 6 | -19.609 | 0.02903 | -0.006554 |
| 2024_n7-5 | 2022_3-5 | 2022_17-1 | 6 | -81.110 | 0.04777 | -0.006477 |
| 2024_n7-6 | 2022_3-5 | 2022_17-1 | 6 | -20.203 | 0.02882 | -0.006489 |
| 2025_7-1 | 2015_5-3 | 2017_13-2 | 5 | 55.285 | -0.00449 | 0.004883 |
| 2025_7-5 | 2015_5-3 | 2017_13-2 | 5 | 69.027 | 0.00280 | -0.003556 |
| 2025_7-7 | 2015_5-3 | 2017_13-2 | 5 | 20.404 | -0.00961 | -0.004538 |
| 2025_7-10 | 2015_5-3 | 2017_13-2 | 5 | 56.499 | -0.00209 | -0.003926 |
| 2025_7-8 | 2015_5-3 | 2017_13-2 | 5 | 46.999 | -0.01751 | -0.000357 |
| 2025_7-9 | 2015_5-3 | 2017_13-2 | 5 | 41.519 | -0.00461 | -0.002823 |
| 2025_7-11 | 2015_5-3 | 2017_13-2 | 5 | 33.838 | 0.00706 | -0.003185 |

| | | | | | | |
|---|---|---|---|---|---|---|
| 2025_5-3 | 2015_16-1 | 2014_4-1 | 6 | 50.882 | 0.01529 | 0.000612 |
| 2025_5-6 | 2015_16-1 | 2014_4-1 | 6 | 20.417 | 0.02001 | -0.004163 |
| 2025_5-13 | 2015_16-1 | 2014_4-1 | 6 | -50.425 | 0.04344 | -0.009502 |
| 2025_12-5 | 2015_6-1 | 2015_6-2 | 3 | -44.321 | -0.00374 | -0.002588 |
| 2025_12-6 | 2015_6-1 | 2015_6-2 | 3 | -16.387 | -0.01003 | -0.004571 |
| 2025_12-10 | 2015_6-1 | 2015_6-2 | 3 | -19.404 | 0.00202 | -0.006799 |
| 2025_4-9 | 2015_12-1 | 2016_16-5 | 6 | 70.829 | 0.00011 | -0.008566 |
| 2025_4-11 | 2015_12-1 | 2016_16-5 | 6 | 0.134 | 0.02165 | -0.010368 |
| 2025_4-12 | 2015_12-1 | 2016_16-5 | 6 | 47.552 | -0.00445 | -0.010590 |
| 2025_4-13 | 2015_12-1 | 2016_16-5 | 6 | -37.215 | 0.03028 | -0.008946 |
| 2025_n8-3 | unknown2 | 2020_5-2 | NA | -2.366 | -0.00588 | -0.001434 |
| 2025_n8-2 | unknown2 | 2020_5-2 | NA | -20.376 | -0.01195 | 0.000700 |
| 2025_n8-4 | unknown2 | 2020_5-2 | NA | -4.570 | -0.00518 | -0.001587 |
| 2025_n3-9 | 2022_2-3 | 2022_4-1 | NA | 57.007 | 0.01429 | -0.003658 |
| 2025_n3-10 | 2022_2-3 | 2022_4-1 | NA | 67.270 | 0.01807 | -0.001409 |
| 2025_n3-14 | 2022_2-3 | 2022_4-1 | NA | 67.488 | 0.05731 | 0.001030 |
| 2025_15-1 | 2024_6-7 | 2024_n7-5 | 7 | -4.412 | 0.03538 | -0.002430 |
| 2025_s1-1 | 2023_9-6 | 2023_15-4 | 3 | 52.915 | -0.00211 | 0.008705 |
| 2025_s1-6 | 2023_9-6 | 2023_15-4 | 3 | 9.995 | 0.00044 | 0.001910 |
| 2025_s1-10 | 2023_9-6 | 2023_15-4 | 3 | 17.713 | 0.00053 | 0.001593 |
| 2025_2-6 | founder_15 | founder_7 | 2 | 65.311 | -0.00326 | 0.000040 |
| 2025_8-7 | founder_17 | founder_12 | 2 | 112.526 | 0.00071 | 0.005116 |
